# Systematic discovery of TIR-based immune signaling systems in bacteria

**DOI:** 10.64898/2025.12.03.692087

**Authors:** Erez Yirmiya, Azita Leavitt, Bohdana Hurieva, Alla H. Falkovich, Nathalie Béchon, François Rousset, Ilya Osterman, Rotem Sorek

## Abstract

Toll/interleukin-1 receptor (TIR) domains are important for immune signaling across humans, plants and bacteria. These domains were recently found to produce immune signaling molecules in plant immunity as well as in a family of bacterial defense systems called Thoeris. Here, we systematically scanned bacterial defense islands to identify anti-phage defense systems involving TIR-mediated signaling. We detected numerous configurations of such systems in bacterial genomes, involving ∼30 different protein effectors predicted to respond to TIR-produced immune signals. We experimentally verified 15 new TIR-containing systems, showing that they provide defense against phages through effector protein domains not previously known as associated with immunity. Further biochemical analyses revealed bacterial Thoeris systems that generate 2′cADPR, an immune signaling molecule central to plant immunity. We also discover multiple types of Thoeris that drive antiphage defense via canonical cADPR, a signaling molecule known to mediate human innate immunity. Our studies show that TIR-based immune signaling systems exist in at least 8% of bacterial genomes, and suggest conservation of TIR-derived immune signals across the tree of life.

## Introduction

Proteins with Toll/interleukin-1 receptor (TIR) domains participate in innate immune recognition and signaling in all domains of life^1^. First identified within animal Toll-like receptors and interleukin receptors, TIR domains have been originally described as responsible for recruitment of downstream immune factors via protein-protein interactions^2,3^. However, recent studies in plants and bacteria showed that TIR domains are enzymes which, once a pathogen is recognized, produce a small signaling molecule that activates downstream immune proteins^4^. In bacteria, TIR immune signaling is carried out by a family of defense systems called Thoeris. Systems from this family employ two core components: a protein containing a TIR domain that senses phage invasion and responds by synthesizing immune signals that are derivatives of the molecule ADP-ribose (ADPR); and an effector protein that is activated by the cognate immune signal to execute premature cell death or cell growth arrest^5,6^.

Three types of Thoeris defense systems were characterized to date in bacteria, all relying on TIR-produced signaling molecules (Figure 1A). In type I Thoeris, the TIR generates 1′′–3′ glycocyclic ADPR (3′cADPR for short), a molecule in which the two ribose moieties in ADPR are linked to each other^7,8^. This molecule binds a SLOG domain in the cognate effector, activating its sirtuin (SIR2) domain to deplete NAD^+^ from the cell and arrest phage replication^6^. In type II Thoeris, the signaling molecule is formed by conjugating ADPR to the amino acid histidine, generating a His-ADPR linear signaling molecule^9^. His-ADPR binds a Macro domain in the effector protein of type II Thoeris, and activates it to form a cell-killing membrane pore^9^. TIR proteins in type IV Thoeris produce N7-cADPR, a variant of cyclic ADPR in which the ribose is covalently attached to the N7 nitrogen atom in the adenine base. A caspase-like protease serves as the effector of type IV Thoeris systems, cleaving cellular proteins indiscriminately when activated by N7-cADPR^10^.

**Figure 1.**
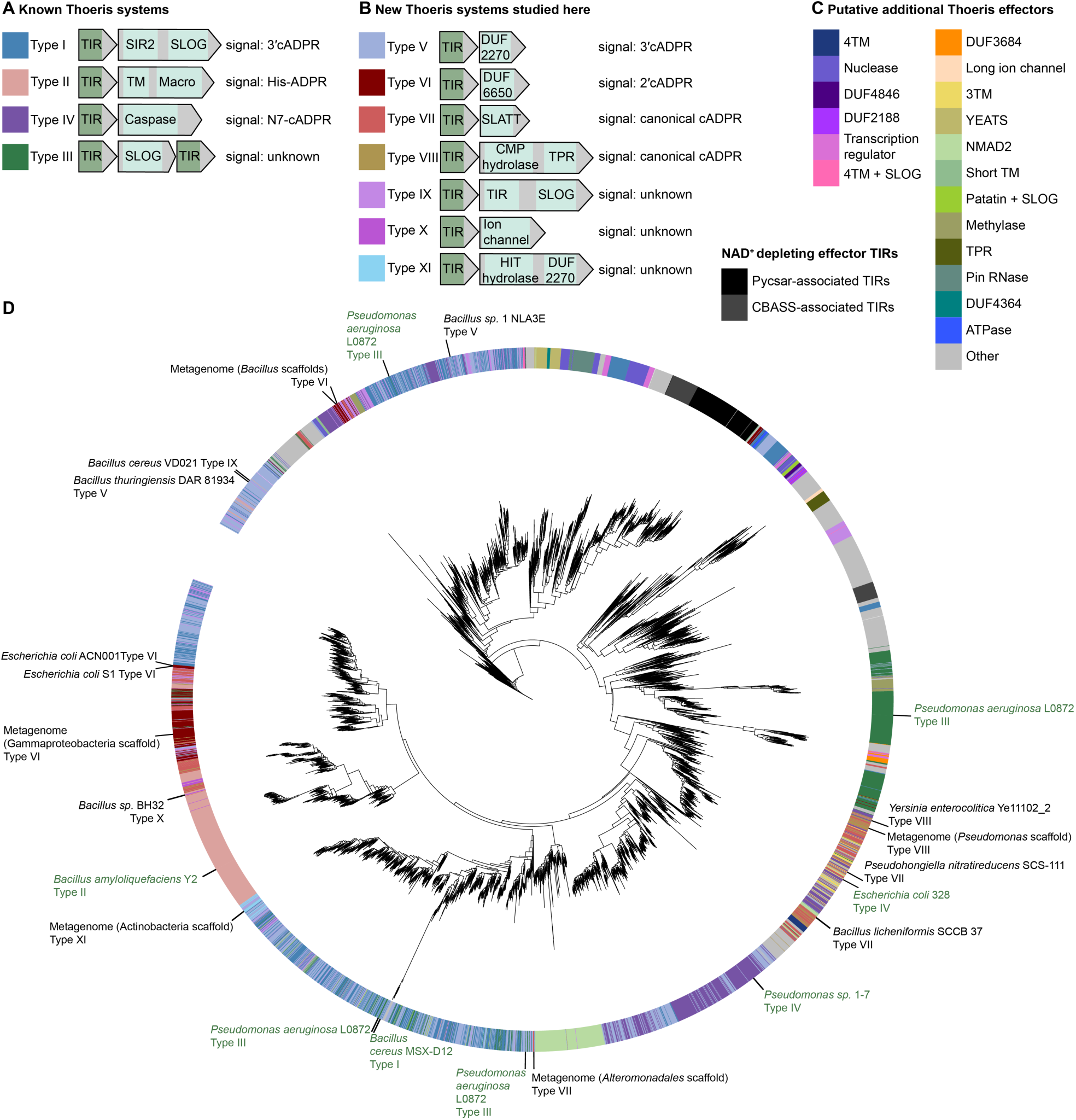
Known and new TIR immune signaling systems in bacteria. (**A**) Known Thoeris systems and their signaling molecules. (**B**) New Thoeris systems discovered here. (**C**) Additional putative effectors frequently found next to TIR-domain proteins in bacteria. (**D**) A phylogenetic tree representing 10,332 TIR domains from TIR-encoding genes that putatively participate in Thoeris immune signaling systems. Systems verified as defensive in this study, and their organism of origin, are indicated on the tree periphery in black. Systems characterized in previous studies are indicated on the tree periphery in dark green. Colors on the outer ring represent the effector associated with the TIR, per the color code shown in panels A-C. The tree also includes, for reference, 604 TIRs known to function as NAD^+^-depleting effectors in CBASS (dark grey) and Pycsar (black) systems.

Understanding TIR signaling in bacteria has been beneficial for deciphering TIR immune signaling in higher organisms, and especially in plants^1,7,11^. Plant TIRs produce signaling molecules that are similar but not identical to the bacterial ones. Notably, a central TIR-produced signal in plants is 1′′–2′ gcADPR (2′cADPR), a molecule similar to the bacterial 3′cADPR, but in which the two ribose moieties are linked via 1′′–2′ O-glycosidic bond rather than 1′′–3′ as in the bacterial immune molecule^7,11,12^. The evolutionary origin of the plant immune signal is currently unknown.

A recent study reported a system called type III Thoeris, which contains multiple TIR-domain proteins coupled with a SLOG domain-containing effector, whose mode of signaling remained uncharacterized^13^. This highlights that despite their importance for bacterial defenses and for understanding immunity in eukaryotes, our understanding of TIR-mediated signaling systems in bacteria is incomplete^14^. In the current study we systematically probed ∼600 million genes from microbial genomes and metagenomes to predict defense systems in which TIRs participate in immune signaling. Our analyses revealed numerous new configurations of candidate Thoeris systems, and detected ∼30 effector proteins predicted to respond to TIR-produced signals. Experimental examination of a subset of these systems verified 15 predicted operons as having anti-phage activities, and validated the production of signaling molecules by the TIR-encoding proteins in response to phage. Some of these bacterial systems were shown to rely on signaling molecules known to mediate plant and animal immune signaling, suggesting that bacterial immune systems formed the ancient origin of eukaryotic TIR signaling.

## Results

### Extensive diversity of Thoeris defense systems in prokaryotes

To comprehensively define the repertoire of Thoeris antiphage defense systems, we analyzed ∼135 million bacterial proteins from ∼38,000 bacterial and archaeal genomes^15^. As sequenced genomes are biased for overrepresentation of species from the *Pseudomonadota* and *Bacillota* phyla, we added ∼500 million proteins from high quality assembled metagenomic scaffolds, to allow for the discovery of Thoeris systems underrepresented in sequenced genomes (Figure S1). All ∼600 million proteins were clustered based on sequence similarity (Methods), and each cluster was analyzed for the tendency of its genes to co-localize with known defense genes in the genomic vicinity, a characteristic indicative of defensive function^5,15–17^. Overall, 341 clusters of proteins with TIR domains, containing a total of 81,720 non-identical proteins, were found to be enriched next to known defense genes. For each of these clusters we analyzed the genomic neighborhood to identify cases in which the genes of the cluster are part of a multigene system^5,15^ (Figure S1).

Some bacterial TIR-domain proteins are known to function as effectors in non-Thoeris defense systems, in which case the TIR depletes NAD^+^ from the infected cells rather than producing a signaling molecule. We discarded TIRs that likely function as NAD^+^-depleting effectors based on their genomic context and neighboring genes. Specifically, TIR domains found in CBASS^18,19^, Pycsar^20^, and TIR-pAgo contexts^21,22^ were discarded. We also removed TIRs in AVAST (Avs)^23,24^ proteins and TIRs found in standalone proteins not associated with other genes in a multigene operon.

We next examined the genes immediately adjacent to the TIR-domain genes in each cluster. In many cases, a substantial fraction of the TIR-domain genes in a given cluster were associated with a known Thoeris effector, but other genes in the cluster were associated with genes not previously described as Thoeris effectors. For example, in one of the clusters there were 302 TIR-domain genes associated with a SIR2-SLOG protein (type I Thoeris), and 160 genes in a type IV Thoeris configuration (Table S1 ; cluster “2612357774”). However, in addition to these, there were 122 TIRs associated with a protein annotated as a domain of unknown function (pfam PF10028, DUF2270) (Table S1). Similarly, a cluster that included 209 cases of TIR genes next to Macro-domain genes (type II Thoeris) also contained 84 TIR genes found next to SLATT-domain genes, and 19 cases of TIRs found next to a predicted ion channel gene (Table S1 ; cluster “2598792572”). We hypothesized that these TIR-containing two-gene operons represent previously undescribed Thoeris systems, and that the associated genes encode effectors that would respond to a signal produced by the TIR-domain protein. Overall, we collected >10,000 non-identical TIR-containing proteins that were deemed as likely representing Thoeris systems (Table S1).

We annotated 29 putative types of Thoeris systems, each with a different predicted effector (Figure 1B, C). In addition to effectors of the four known Thoeris systems, putative effectors included various predicted enzymes such as DNases, RNases, phospholipases, and hydrolases, as well as a variety of proteins with transmembrane (TM)-spanning helices and proteins with domains of unknown function (Figure 1). Some of these domains were previously found in cell-killing effectors of CBASS and other defense systems, supporting their predicted roles as effectors in the Thoeris systems predicted here^19,25–27^.

To assess whether the predicted operons can defend against phages, we selected 10 new configurations of putative Thoeris systems for experimental investigation. Several representatives were chosen for each configuration, mostly selecting source organisms closely related taxonomically to *Escherichia coli* or *Bacillus subtilis* to increase chances for compatibility when heterologously expressing the tested systems in these lab bacteria. Overall, 30 TIR-containing operons were cloned either in *E. coli* MG1655 or in *B. subtilis* BEST7003 strains (Table S2). These bacteria were then challenged with a diverse set of 22 phages (12 *E. coli* phages and 10 *B. subtilis* phages), leading to the verification of 15 of the tested systems as able to defend against phages (Figure 2, Figure S2).

**Figure 2.**
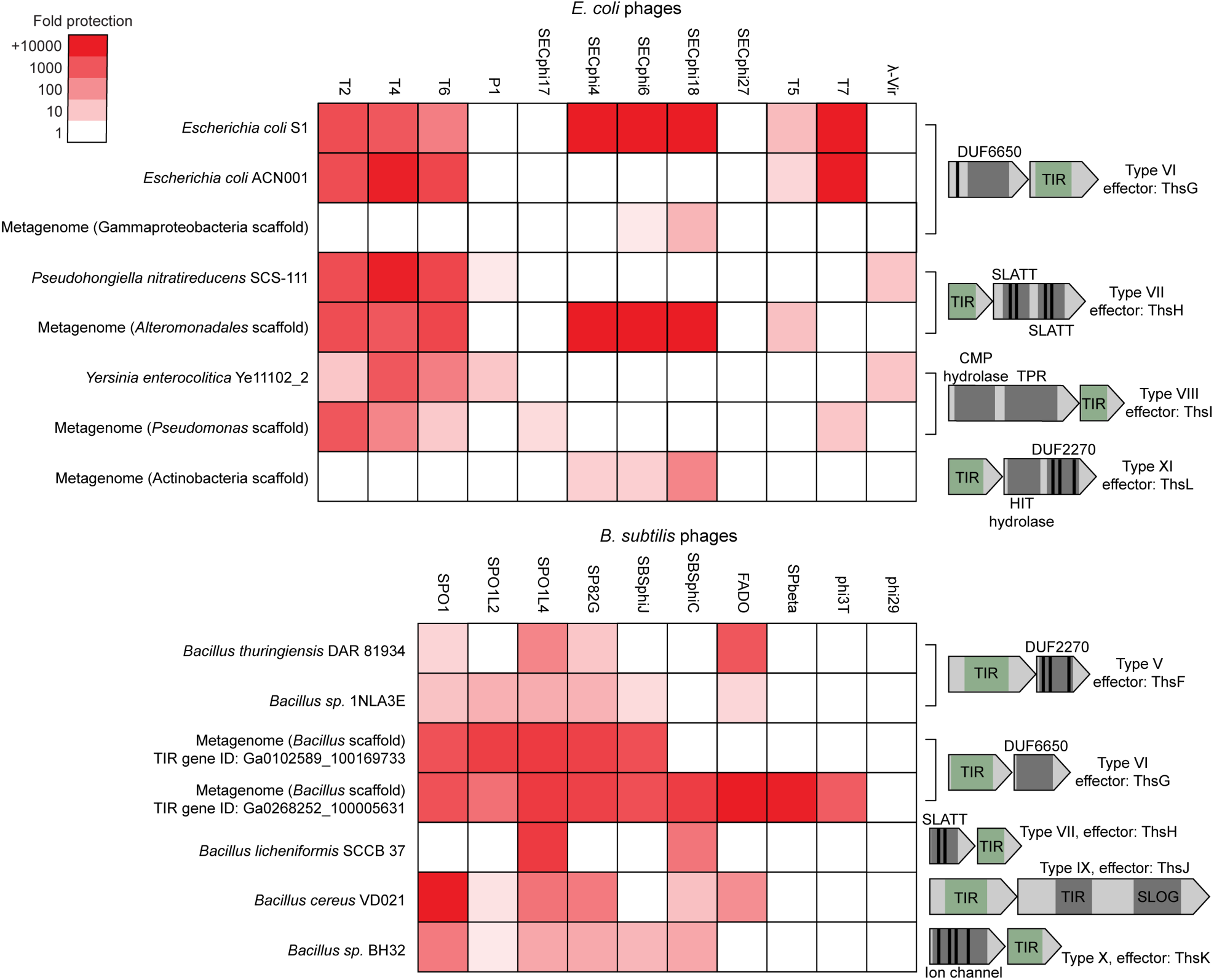
New Thoeris systems verified to protect against phages. Heatmap showing fold antiphage protection conferred by Thoeris systems cloned and genomically integrated (except for the type XI system) into *E. coli* (top) or *B. subtilis* (bottom). Protection levels were determined by serial dilution plaque assays, comparing strains harboring the indicated Thoeris system with control strains carrying an empty vector. Data represent the mean of three independent replicates; individual datapoints are shown in Figure S2. On the right, schematic diagrams illustrate the genetic organization of the validated Thoeris types, with signaling TIR domains in green, predicted domains in the effector in grey, and transmembrane helices in black.

The defensive operons validated seven new configurations of Thoeris systems, which we henceforth name Thoeris types V, VI, VII, VIII, IX, X and XI (Figures 1, 2). We expand the gene nomenclature originally introduced for type I Thoeris, where the TIR domain gene was named *thsB* and the effector *thsA*^6^, and name effectors of other Thoeris types in an alphabetical order. Hence, the effectors of the previously described Thoeris types are now called ThsA, ThsC, ThsD and ThsE for types I, II, III and IV respectively; with ThsF, ThsG, ThsH, ThsI, ThsJ, ThsK and ThsL standing for effector proteins of new Thoeris types (Figure 1, 2).

### Type V Thoeris employs 3′cADPR to trigger DUF2270 effector toxicity

We next sought to characterize the immune signaling molecules produced by the various new Thoeris types. It was previously shown that when a Thoeris TIR is expressed alone in a bacterial cell, without the cognate effector, it can be triggered by phage infection to produce substantial amounts of the signaling molecule which accumulates in the infected cell^6,7,10,28^. We therefore prepared cells expressing only the TIR proteins from the 15 verified systems, infected each of these cells by a phage that is naturally blocked by the respective defense system, and collected filtered lysates from the infected cells. We hypothesized that these lysates contain the signaling molecule produced by the expressed TIR during phage infection (Figure 3A).

**Figure 3.**
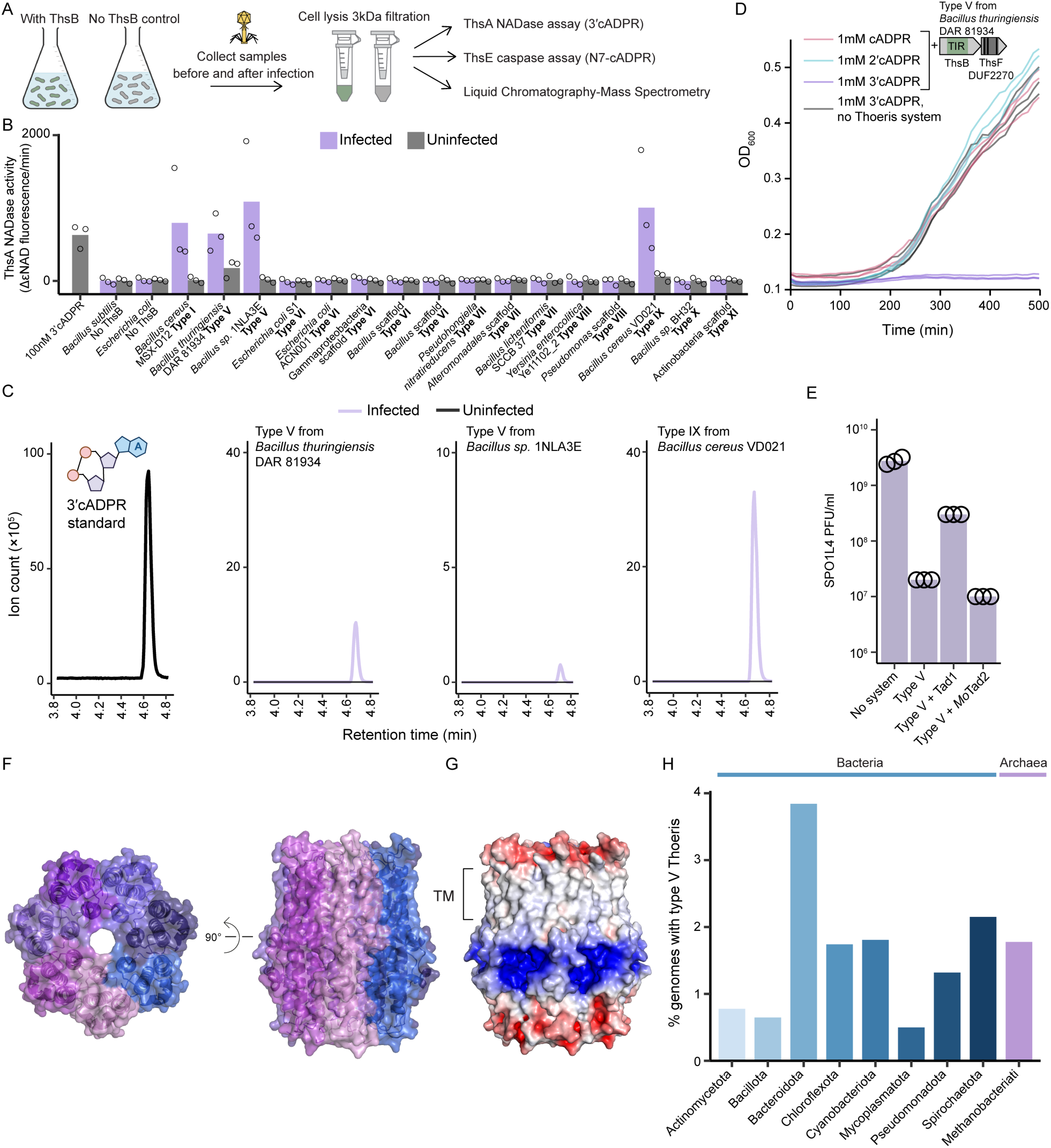
ThsB of type V Thoeris employs 3′cADPR to activate DUF2270 effector toxicity. (**A**) Assays applied to identify signaling molecules produced by ThsB proteins from Thoeris types V-XI. (**B**) Cells expressing ThsB from the indicated Thoeris systems were infected with phage to trigger ThsB immune signal production, and then lysed and filtered to enrich for small molecules. NADase activity of purified ThsA incubated with the filtered lysates was measured using a nicotinamide 1,N6-ethenoadenine dinucleotide (εNAD) cleavage fluorescence assay. Bars represent the means of three experiments, with individual data points overlaid. (**C**) Extracted ion chromatograms (m/z = 540.053) from lysates of cells expressing the TIR domain of the indicated system, before and after infection. Left: chromatogram of a 3′cADPR standard, with the structure of the molecule shown above. Data are representatives of three replicates. (**D**) Growth curves of *B. subtilis* cells expressing the type V Thoeris system from *B. thuringiensis* DAR 81934 or an empty vector, supplemented with different cyclic ADPR isomers. Three repeats are shown for each condition. (**E**) Plaque assay results of phage SPO1L4 infecting *B. subtilis* cells expressing the type V Thoeris system from *B. thuringiensis* DAR 81934, both the type V Thoeris system and anti-Thoeris sponge proteins, or an empty vector. Data represent PFUs per milliliter, and bars show the average of three repeats with individual data points overlaid. (**F**) Alphafold3 model of the type V Thoeris effector from *B. thuringiensis* DAR 81934 as a homo-heptamer (ipTM score = 0.78). (**G**) Electrostatic potential map of the effector presented in panel F (negative charge, red; positive, blue). (**H**) Detection of type V Thoeris in prokaryotic genomes, shown for phyla with at least 200 genomes in the downloaded database. The data for “no Thoeris system” control presented in panel 3D is also presented in figures S3, S4A, and S4B. The data for the type V Thoeris system from *B. thuringiensis* DAR 81934 presented in panels 3E and S2A are the same.

To ask whether any of the TIRs produce the same signaling molecule as type I Thoeris (3′cADPR), we took advantage of a previously established biochemical assay that allows sensitive detection of 3′cADPR^6,7^. This assay is based on in vitro activation of *B. cereus* ThsA, an enzyme activated by 3′cADPR but not by closely related molecules such as 2′cADPR or any other known Thoeris signal^7,11^. Filtered lysates from cells expressing the TIR proteins of types V and IX Thoeris were able to activate ThsA in vitro, suggesting that these TIRs produce 3′cADPR (Figure 3B). ThsA activation was minimal when lysates were taken from TIR-expressing uninfected cells, verifying that the TIRs produce the signal in response to phage (Figure 3B). Liquid chromatography and mass spectrometry (LC-MS) confirmed infection-dependent 3′cADPR synthesis in these lysates, establishing that the TIR proteins from Thoeris types V and IX generate 3′cADPR upon phage detection (Figure 3C).

Previous studies have shown that the signaling molecules of signaling systems such as Pycsar can penetrate bacterial cells when added at high-micromolar concentrations to the growth medium and can trigger cognate effector toxicity^20,29^. We verified that addition of synthetic 3′cADPR to growth media of cells expressing type I Thoeris causes Thoeris-mediated toxicity in the absence of phage infection, showing that this molecule can penetrate cells as well (Figure S3). Growth of cultures expressing the type V Thoeris from *B. thuringiensis* DAR 81934 was impaired when 3′cADPR was supplemented to the growth media, suggesting that the effector protein in this Thoeris system is triggered by 3′cADPR (Figure 3D). Other cADPR isomers, including 2′cADPR and canonical cADPR, were not toxic to cells expressing the type V Thoeris from *B. thuringiensis*, demonstrating the specificity of this system to 3′cADPR (Figure 3D). 3′cADPR was not toxic to cells expressing another variant of type V Thoeris from *Bacillus* sp 1. NLA3E, nor was it toxic to cells expressing type IX Thoeris, implying that the effectors in these systems may be activated by another molecule or require an additional infection-derived signal to be activated (Figure S4; See Discussion).

Phages can evade Thoeris defense by producing sponge proteins that sequester the immune signaling molecule produced by the TIR^7,9,28–30^. Co-expression of the type V system from *B. thuringiensis* with Tad1 from phage SBSphiJ7, a sponge protein that specifically sequesters 3′cADPR, resulted in substantially reduced defense, providing further support that the *B. thuringiensis* type V system relies on 3′cADPR as an immune signaling molecule. In contrast, *Mo*Tad2, a sponge that sequesters the Thoeris type II signal His-ADPR but is incapable of sequestering 3′cADPR^9^, did not inhibit defense by type V Thoeris (Figure 3E). Together, these results establish 3′cADPR as the signaling molecule of the *B. thuringiensis* DAR 81934 type V Thoeris system.

The effector of type V Thoeris is predicted to encode three membrane-spanning helices and is hence likely to function via a membrane-targeting mechanism. This effector is annotated as belonging to a protein family of unknown function (pfam: PF10028, DUF2270), which we now rename ThsF. Structural modeling using AlphaFold3 (AF3) revealed a conserved, circular homo-oligomeric assembly generating a channel-like structure that is likely membrane-spanning (Figures 2, 3F,G; Figure S5). Within the predicted homo-oligomeric structure of ThsF, deep, positively charged pockets are formed at the interfaces between protomer pairs (Figure 3G). Presumably, 3′cADPR binding in these pockets induces conformational changes that activate membrane disruption, thereby leading to toxicity and interrupting phage replication. Notably, binding sites for 3′cADPR and other nucleotide-containing immune signaling molecules are typically positively charged to accommodate the negative charge of the immune signal^30^. We detected 470 type V Thoeris systems in about 1.2% of the analyzed genomes, a fraction comparable to the fraction of genomes harboring the previously known Thoeris types (Table S3). Type V systems are present in genomes of diverse phyla and are especially enriched in *Bacteroidota* (Figure 3H).

### Type VI Thoeris utilizes 2′cADPR, revealing signaling conservation with plant immunity

We next asked whether any of the TIRs from the other Thoeris systems produces the signaling molecules known for type II Thoeris (His-ADPR) or type IV Thoeris (N7-cADPR). Lysates from infected TIR-expressing cells did not activate the caspase-like effector of type IV Thoeris in vitro^10^, suggesting that none of the new Thoeris systems produces N7-cADPR in response to infection. As there is currently no enzyme-based in vitro assay to measure His-ADPR, we attempted to identify His-ADPR in lysates of infected cells that express the new TIRs via LC-MS, but none of these lysates exhibited detectable amounts of this molecule (Figure S6). These results suggest that the 12 remaining new Thoeris systems (those that were not found to produce 3′cADPR) generate a signaling molecule not previously reported in bacterial immunity.

We reasoned that some Thoeris systems might produce the same signaling molecules as those generated by plant immune TIRs. We specifically focused on 2′cADPR, an immune signal broadly synthesized by plant TIRs in response to pathogens^7,8,11,12,31^. To search for bacterial TIRs that may produce 2′cADPR, we developed an LC-MS protocol that allows separation of cADPR isomers (Figure S7), and used this protocol to examine lysates from infected TIR-expressing cells. Three TIR proteins in our set were found to produce substantial amounts of 2′cADPR in response to phage, but not in the absence of infection (Figures 4A). Remarkably, the effectors associated with all three TIRs were similarly annotated as proteins with a domain of unknown function DUF6650 (pfam PF20355). We denote the TIR-DUF6650 systems as type VI Thoeris, with the effector protein named ThsG (Figure 4).

**Figure 4.**
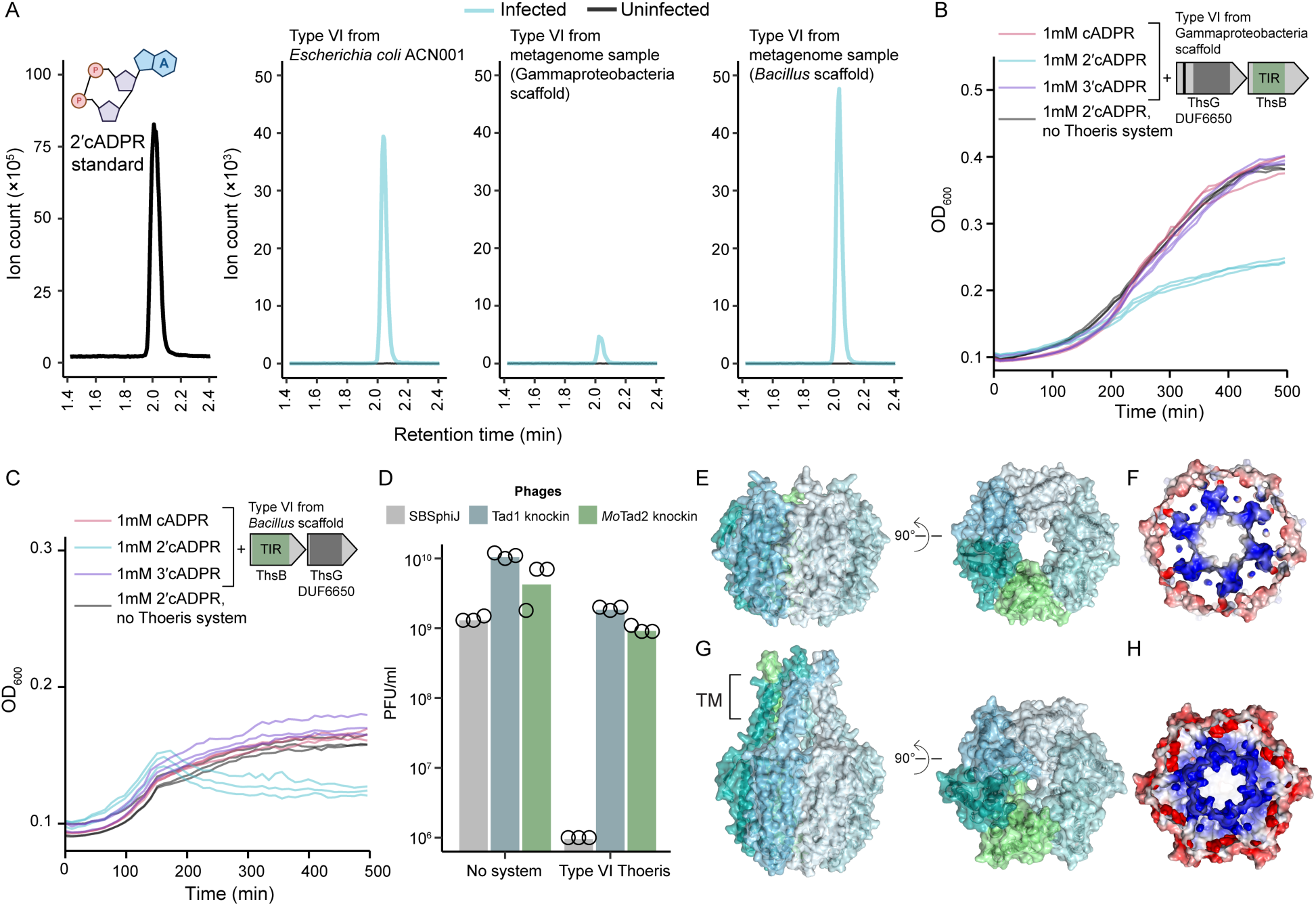
Type VI Thoeris utilizes 2′cADPR as an immune signal. (**A**) Extracted ion chromatograms (m/z = 540.053) from lysates of cells expressing the TIR domain of the indicated system, before and after infection. Left: chromatogram of a 2′cADPR standard, with the structure of the molecule shown above. Data are representatives of three replicates. (**B**) Growth curves of *E. coli* cells expressing the type VI Thoeris system from the metagenomic Gammaproteobacteria scaffold or an empty vector. Bacteria were grown in minimal media (Methods), and media were supplemented with the indicated cyclic ADPR isomers. Three repeats are shown as individual curves for each condition. (**C**) Growth curves of *B. subtilis* cells expressing the type VI Thoeris system from the metagenomic *Bacillus* scaffold or an empty vector. Bacteria were grown in minimal media (Methods), and media were supplemented with the indicated cyclic ADPR isomers. Three repeats are shown for each condition. (**D**) Plaque assay results of phage SBSphiJ, SBSphiJ with a knocked-in Tad1, and SBSphiJ with a knocked-in *Mo*Tad2, infecting *B. subtilis* cells expressing the type VI Thoeris system from the metagenomic *Bacillus* scaffold or an empty vector. Data represent PFUs per milliliter, and bars show the average of three repeats with individual data points overlaid. (**E**) Alphafold3 model of the type VI Thoeris effector from the metagenomic *Bacillus* scaffold as a homo-hexamer (ipTM score = 0.91). (**F**) Electrostatic potential map of the effector presented in panel E (negative charge, red; positive, blue). (**G**) Model of the type VI Thoeris effector from the metagenomic Gammaproteobacteria scaffold as a homo-hexamer (ipTM score = 0.79). (**H**) Electrostatic potential map of the effector in panel G. The data for phage SBSphiJ infecting negative control cells and cells expressing the type VI Thoeris system from the metagenomic *Bacillus* scaffold presented in panel 4D is also presented in figure S2A.

Supplementing 2′cADPR to the growth medium induced growth arrest in cells harboring type VI systems, providing support that 2′cADPR functions as the cognate immune signal in these systems (Figure 4B-C). Other cADPR isomers, including 3′cADPR and canonical cADPR, did not affect the growth of these cells, further demonstrating the specificity of the type VI system to 2′cADPR (Figure 4B,C). 2′cADPR inhibited growth of *E. coli* cells expressing a type VI system taken from a metagenomic scaffold of Gammaproteobacteria origin, and of *B. subtilis* cells expressing a type VI system taken from a metagenomic scaffold of *Bacillus* origin, confirming that 2′cADPR is the immune signal of type VI Thoeris systems from distantly related bacteria (Figure 4B,C).

We have previously shown that Tad1 from phage SBSphiJ7 is able to efficiently sequester 2′cADPR in vitro^7,28^. Incubation of purified *Mo*Tad2 with 2′cADPR demonstrated that *Mo*Tad2 also sequesters this molecule, as confirmed by high-performance liquid chromatography (HPLC) (Figure S8). We therefore tested whether phages engineered to express either Tad1 or *Mo*Tad2 can overcome type VI Thoeris defense. While phage SBSphiJ was blocked when infecting *B. subtilis* cells expressing type VI Thoeris, SBSphiJ strains knocked-in for Tad1 or *Mo*Tad2 successfully infected cells even when those expressed type VI Thoeris (Figure 4D). Together, these findings establish that type VI Thoeris systems utilize 2′cADPR for immune signaling, revealing conservation of immune signaling molecules between bacterial and plant TIR-dependent defense systems.

Type VI Thoeris systems are considerably rarer in nature than type V systems. We detected 255 type VI Thoeris operons in microbial genomes (Table S3). Most of the systems were found in species belonging to the bacterial phyla *Pseudomonadota*, *Bacillota, Actinomycetota,* and *Cyanobacteriota*, although rare appearances were recorded also in *Bacteroidota*, *Spirochaetota*, and other phyla (Table S3). We found only two type VI systems in archaea.

The ThsG effector of type VI Thoeris is not similar to any domain of known function. AF3 consistently models this effector and its homologs as a homo-hexamer with high confidence (Figure 4E,F; Figure S9). This complex forms a tube-shaped circular structure with positively charged pockets formed between protomers on the inner lumen of the tube (Figure 4E,F). In some cases, the ThsG effector contains N-terminal transmembrane-spanning helices in addition to the DUF6650 domain (Figure 4G,H; Figure S9). In these cases, 2′cADPR binding may activate the protein to breach membrane integrity and halt phage propagation. It is currently unclear how ThsG effectors that lack transmembrane-spanning helices can exert toxicity.

### Canonical cADPR functions as an immune signal in bacterial immunity

We next sought to characterize the immune signaling molecules produced by the TIRs of the remaining new Thoeris systems. Nine TIRs in our set were found to produce neither 3′cADPR nor 2′cADPR. However, for 5 of these TIRs the LC-MS data showed the appearance of a mass with both an m/z value and retention time equivalent to the canonical cADPR molecule, suggesting that these TIRs produce cADPR as their signaling molecule (Figures 5A , S7). Production of cADPR by these TIRs was strictly phage-dependent, with no detection in uninfected controls (Figure 5A). Three of the TIRs found to produce cADPR were associated with an effector having a SLATT domain annotation, and two additional TIRs were associated with an effector annotated as a CMP hydrolase (Figure 2). We denote these two configurations as types VII and VIII Thoeris, and the effectors proteins as ThsH and ThsI, respectively.

**Figure 5.**
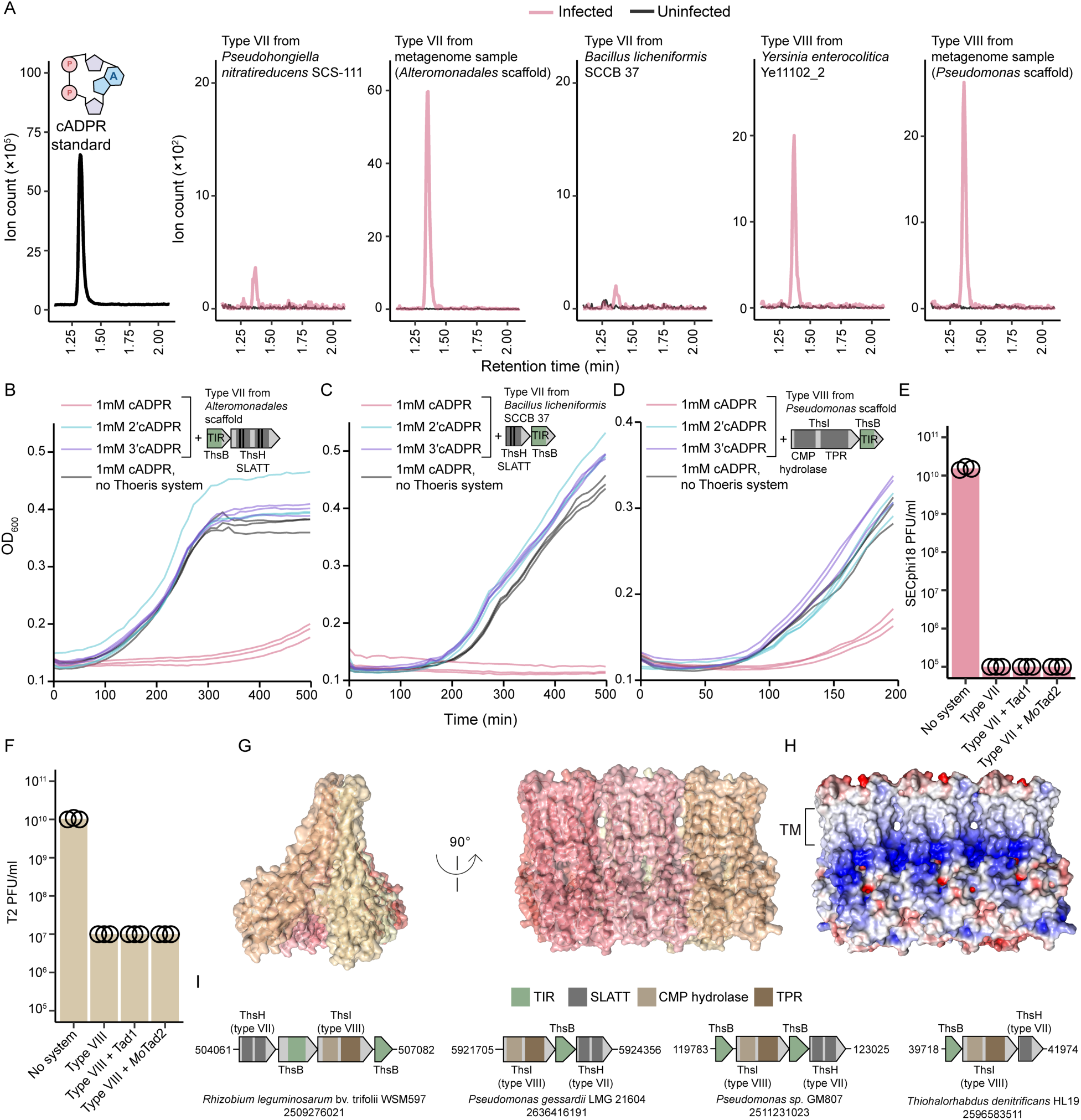
Canonical cADPR acts as the immune signal of types VII and VIII Thoeris. **(A)** Extracted ion chromatograms (m/z = 540.053) from lysates of cells expressing the TIR domain of the indicated system, before and after infection. Left: chromatogram of a canonical cADPR standard, with the structure of the molecule shown above. Data are representatives of three replicates. (**B**) Growth curves of *E. coli* cells expressing the type VII Thoeris system from the metagenomic *Alteromonadales* scaffold or an empty vector. Medium was supplemented with the indicated cyclic ADPR isomers. Bacteria were grown in room temperature (RT). (**C**) Growth curves of *B. subtilis* cells expressing the type VII Thoeris system from *Bacillus licheniformis* SCCB 37 or an empty vector; medium was supplemented with cyclic ADPR isomers. Bacteria were grown in RT. (**D**) Growth curves of *E. coli* cells expressing the type VIII Thoeris system from the metagenomic *Pseudomonas* scaffold or an empty vector; medium was supplemented with cyclic ADPR isomers. Bacteria were grown at 37 °C. For panels B, C and D, three repeats for each condition are shown as individual curves. (**E**) Plaque assay results of phage SECphi18 infecting *E. coli* cells expressing the type VII Thoeris system from the metagenomic *Alteromonadales* scaffold, or coexpressing the type VII Thoeris system with Tad1 or *Mo*Tad2. Bacteria with an empty vector (No system) was used as control. (**F**) Plaque assay results of phage T2 infecting *E. coli* cells expressing the type VIII Thoeris system from the metagenomic *Pseudomonas* scaffold, or coexpressing the type VII Thoeris system with Tad1 or *Mo*Tad2. Bacteria with an empty vector (No system) was used as control. For panels E and F, Data represent PFUs per milliliter, and bars show the average of three repeats with individual data points overlaid. (**G**) Alphafold3 model of the type VII Thoeris effector from the metagenomic *Alteromonadales* scaffold as a homo-hexamer (ipTM score = 0.76). (**H**) Electrostatic potential map of the model shown in panel G (negative charge, red; positive, blue). (**I**) Genomic loci in which both ThsH and ThsI effectors are encoded in the same Thoeris operon. Source organism and IMG taxon IDs are indicated below each operon. The data for the type VII Thoeris system from the metagenomic *Alteromonadales* scaffold presented in panels 5E and S2D are the same. The data for the type VIII Thoeris system from the metagenomic *Pseudomonas* scaffold presented in panels 5F and S2E are the same.

Bacteria expressing types VII and VIII Thoeris were unable to grow when cADPR was supplemented to their growth media, but growth was not impaired in the presence of 2′cADPR or 3′cADPR, suggesting that cADPR specifically activates the effector in these systems (Figure 5B-D). Toxicity was observed both for systems of Proteobacterial origin expressed in *E. coli* and for a system from *Bacillus licheniformis* expressed in *B. subtilis*, confirming dependence on cADPR signaling for types VII and VIII systems across a range of bacteria (Figure 5B-D). Expression of Tad1 or *Mo*Tad2 together with the defense systems did not affect their ability to defend against phages, further confirming that these systems do not rely on 3′cADPR, 2′cADPR, or His-ADPR for immune signaling (Figure 5E,F).

The canonical cADPR, in which the ribose is bound to the N1 nitrogen atom of the adenine base, is a well-known immune signaling molecule in humans^32–34^. Our finding of cADPR-based immune signaling in bacteria therefore possibly points to ancient evolutionary connections between bacterial antiviral immunity and human immune signaling pathways (see Discussion).

The effector protein in type VII Thoeris, ThsH, is annotated with a SLATT domain. SLATT domains are found in membrane-spanning proteins known to function as cell-killing effectors in CBASS systems, where they are thought to oligomerize into membrane-breaching assemblies when activated by CBASS-produced cyclic dinucleotides^19,35^. It is therefore likely that ThsH effectors similarly oligomerize upon binding to the cADPR molecule produced by the TIR of type VII Thoeris. Indeed, AF3 confidently predicts ThsH to form a filament-like transmembrane-spanning structure, in which the cytoplasmic region contains positively charged pockets formed between SLATT protomers (Figure 5G,H).

Type VIII Thoeris effectors (ThsI) have a protein domain annotated as CMP hydrolase. This domain is found in families of nucleotide-hydrolyzing enzymes involved in nucleotide and nucleoside biosynthesis and degradation pathways, including enzymes that degrade specific nucleotides or contribute to the biosynthesis of modified nucleosides^36,37^. We therefore hypothesize that once ThsI is activated by cADPR, it will manipulate the nucleotide pool to deny the phage of an essential molecule, as done by many bacterial defense systems^38^. ThsI also contains a tetratricopeptide repeat (TPR) domain, which is a domain known to form a platform for binding other molecules. Presumably, the TPR domain may be responsible for the perception of cADPR by ThsI. Notably, this TPR domain was previously predicted by Aravind and colleagues to participate in bacterial TIR-mediated signaling^39^.

The cADPR-dependent types VII and VIII Thoeris systems are abundant in bacterial genomes and are cumulatively present in about 2% of sequenced genomes. While type VII systems are spread across *Pseudomonadota*, *Actinomycetota*, *Bacillota*, *Cyanobacteriota*, and other phyla, type VIII systems are especially abundant in *Pseudomonadota* (Table S3). In rare occasions we observe Thoeris systems in which both ThsH and ThsI effectors are present in the same TIR-containing operon, consistent with our observation that both respond to the same signaling molecule (Figure 5I).

### Distribution of Thoeris systems in Bacterial and Archaeal genomes

It was previously thought that TIR-mediated signaling is rare in bacteria as compared to CBASS immune signaling^40^. However, our data now show that Thoeris systems, including the four previously known ones and the seven new systems discovered here, are cumulatively present in ∼8% of analyzed bacterial and archaeal genomes (Figure 6A-C, Table S3). If the other Thoeris systems predicted here but not yet verified (Figure 1C) indeed function in immune signaling, we surmise that even a larger fraction of all bacterial and archaeal genomes encode Thoeris, making these systems in the same ballpark of abundance as CBASS and AVAST systems^19,23,24^. Our data now show that immunity based on intracellular signaling with modified nucleotide signals is an extremely successful defense strategy in bacteria: we found that 19.7% of the bacteria and archaea in our set encode at least one of the CBASS, Thoeris, or Pycsar immune signaling systems (Figure 6C, Table S3).

**Figure 6.**
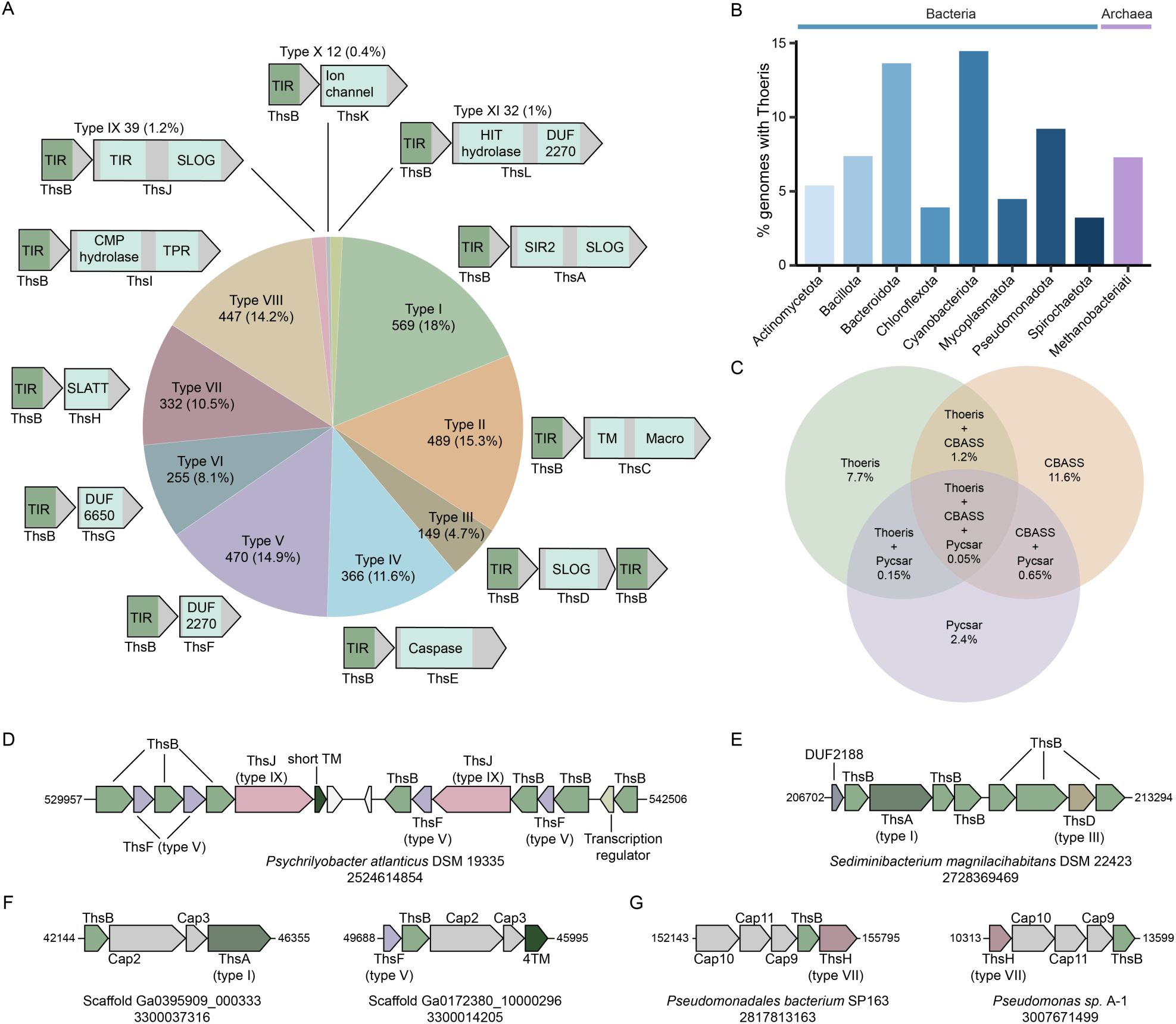
Distribution of Thoeris systems across microbial genomes. (**A**) Distribution of 3,154 detected Thoeris systems in 38,149 bacterial and archaeal genomes. (**B**) Fraction of genomes encoding Thoeris systems across bacterial and archaeal phyla, shown for phyla with at least 200 genomes in the dataset. (**C**) A diagram showing the percentage of genomes encoding Thoeris, CBASS, and Pycsar systems in the downloaded database. Percentages represent the fraction of genomes encoding each system without deduction of overlapping genomes (e.g., 7.7% of genomes encode a Thoeris system regardless of whether they also encode CBASS or Pycsar). **(D,E)** Representative defense islands encoding multiple ThsB and Thoeris effector proteins. The source organism and IMG taxon IDs are indicated below each genomic region. (**F**) Thoeris operons encoding Cap2 and Cap3. Scaffold of origin and metagenomic sample ID are indicated below each operon. (**G**) Thoeris operons encoding Cap9, Cap10, and Cap11. The source organism and IMG taxon ID are indicated below each operon.

Most bacteria with Thoeris encode one such system in their genomes, although we observed rare cases of multiple systems in the same genome. *Psychrilyobacter atlanticus* DSM 19335, for example, encodes seven ThsB proteins, four ThsF proteins (type V Thoeris), two ThsJ proteins (type IX) and two additional putative effectors that we predicted but not experimentally tested, all in the same defense island (Figure 6D). Similarly, *Sediminibacterium magnilacihabitans* DSM 22423 has six ThsB proteins, ThsA (type I), ThsD (type IV), and a putative effector with a domain of unknown function (DUF2188), all encoded in the same genomic region (Figure 6E).

While the core operon of Thoeris generally comprises the TIR domain and the effector genes, some Thoeris operons are more complex. For example, we found Thoeris operons that encode, in addition to the TIR and the effector, also Cap2 and Cap3 proteins (Figure 6F). These proteins were documented as ancillary proteins in type II CBASS systems, where they conjugate the CBASS CD-NTase onto cellular targets via a machinery reminiscent of eukaryotic ubiquitin conjugation^41,42^. TIR proteins operonically associated with *cap2* and *cap3* display the typical C-terminal tail necessary for Cap2/3 mediated conjugation^43^, suggesting that the ancillary proteins in these systems function similar to type II CBASS (Figure S10A). We also found operons similar to type IV CBASS, in which the TIR and effector proteins are associated with *cap9*, *cap10* and *cap11* genes (Figure 6G). AF3 modeling predicts that the TIR in these operons interacts with Cap9 and Cap10 in a manner similar to that shown for the CD-NTase in type IV CBASS^44^, suggesting a similar mechanism (Figure S10B).

## Discussion

Our findings reveal that TIR-based immune signaling is far more common and functionally diverse in bacteria than previously appreciated. By systematically scanning ∼600 million microbial proteins, we mapped dozens of distinct Thoeris architectures and expanded the Thoeris family to eleven experimentally verified system types. These systems collectively appear in ∼8% of bacterial and archaeal genomes, establishing Thoeris as a prevalent antiviral defense strategy in prokaryotes.

The 2′cADPR molecule, originally discovered as produced by the monocot plant *Brachypodium distachyon* TIR-domain protein BdTIR^6,7^, is now recognized as a central immune signaling molecule in plants^11,12,31^. This molecule is produced by plant TIR-domain proteins once they recognize a signature of pathogen infection^1,12,45,46^, and recent evidence suggest that it may be further processed in the plant cell to become phosphoribosyl adenosine monophosphate (pRib-AMP)^12,31^. The plant immune signal is sensed by a protein complex involving the protein EDS1 and other factors, leading to propagation of the plant immune response^45,47^. Until our study, 2′cADPR immune signaling was considered exclusive to plants, although the finding that phage sponge proteins can bind and sequester 2′cADPR hinted that it might have a role in bacterial immunity as well^7,28^. Our finding establishes 2′cADPR as an abundant immune signal produced by diverse bacteria, further strengthening the hypothesis that TIR-mediated immune signaling originated in prokaryotes^48^.

The canonical cADPR, in which ADPR is cyclized via a connection between the terminal ribose and the adenine base, is a molecule long known as associated with muscle, neural and immune functions in animals^49^. This molecule activates calcium channels of the ryanodine receptor family, resulting in Ca^2+^ influx that can regulate diverse functions such as muscle contraction, insulin secretion, neurotransmitter release, immune cell activation and cytokine release^33,49–51^. In humans and other animals, cADPR is produced by the TIR-domain protein SARM1 in the process of injury-induced regulated neuronal axon death, but it is currently unknown whether cADPR is directly involved in regulating the axon death process^52–55^. The membrane-associated protein CD38 in human, which is an ADP-ribosyl cyclase of a protein family unrelated to TIR, also produces cADPR in animals^49^. Based on our findings that cADPR is produced by bacterial TIR proteins and participates in immune signaling, we predict that TIR-dependent cADPR production will be found important for immune signaling in eukaryotes in future studies.

While our study revealed the immune signaling molecules of four new types of Thoeris, the molecules remain unknown for a number of other types. The TIR domains of types X and XI Thoeris (associated with an ion channel and a hydrolase effector, respectively) were not found to produce any of the known TIR-derived molecules in response to infection, suggesting that these TIRs possibly produce new kinds of signaling molecules. In addition, while the TIR of type IX Thoeris was shown to produce 3′cADPR following phage infection, this molecule did not activate effector toxicity of this system, suggesting that another molecule might be the active one in this system, or that its activation necessitates an additional signal on top of 3′cADPR. Together with the predicted Thoeris systems that were not yet studied (Figure 1C), these observations indicate that there are additional TIR-derived immune signals and activation mechanisms remaining to be discovered. Defining these missing molecules and effectors will not only complete the biochemical landscape of Thoeris immunity, but may once again contribute to our understanding of immune signaling in the eukaryotic realm.

### Limitations of the study

While the cluster-based detection of new Thoeris configurations enabled the identification of 29 putative Thoeris types, it should be noted that the effector annotations presented in Figure 1 and Table S1 are based on protein representatives and therefore include a certain degree of noise. These annotations should not be used to calculate the distribution of Thoeris types across genomes. Instead, accurate estimation of type distribution better rely on the detection of Thoeris systems in microbial genomes using DefenseFinder^40^, which was used in the current study to analyze the distribution of the eleven experimentally validated Thoeris types, but not of the remaining putative effectors that were not experimentally studied or verified. Additionally, although TIR proteins from the putative Thoeris types not verified in this work cluster together with TIRs established to function as ThsB signaling proteins, it remains possible that in some of the non-verified putative Thoeris types the TIR domain acts as an effector that depletes NAD⁺ upon activation. Such systems would therefore not represent true Thoeris configurations, but rather immune pathways in which the TIR serves as the toxic effector as seen in other defense systems^19–25^.

Another point worth noticing is that two of the five tested type VI systems did not produce detectable 2′cADPR upon phage infection. A possible explanation is that the production of 2′cADPR by the TIRs in these systems is weak and falls under the limits of detection; but another possibility is that these variants of type VI systems rely on a different immune signal that remains to be identified.

## Acknowledgements

We thank members of the Sorek lab for constructive discussions during this study. R.S. was supported, in part, by the European Research Council (grant ERC-AdG GA 101018520), the Israel Science Foundation (MAPATS grant 2720/22), the Deutsche Forschungsgemeinschaft (SPP 2330, grant 464312965), the Minerva Foundation with funding from the Federal German Ministry for Education and Research, a research grant from Magnus Konow in honor of his mother Olga Konow Rappaport, the Center for Immunotherapy at the Weizmann Institute of Science, and the Knell Family Center for Microbiology. I.O. was supported by the Ministry of Absorption New Immigrant program. E.Y. was supported by the Clore Scholars Program and, in part, by the Israeli Council for Higher Education (CHE) via the Weizmann Data Science Research Center.

## Supplementary figures

**Figure S1.**
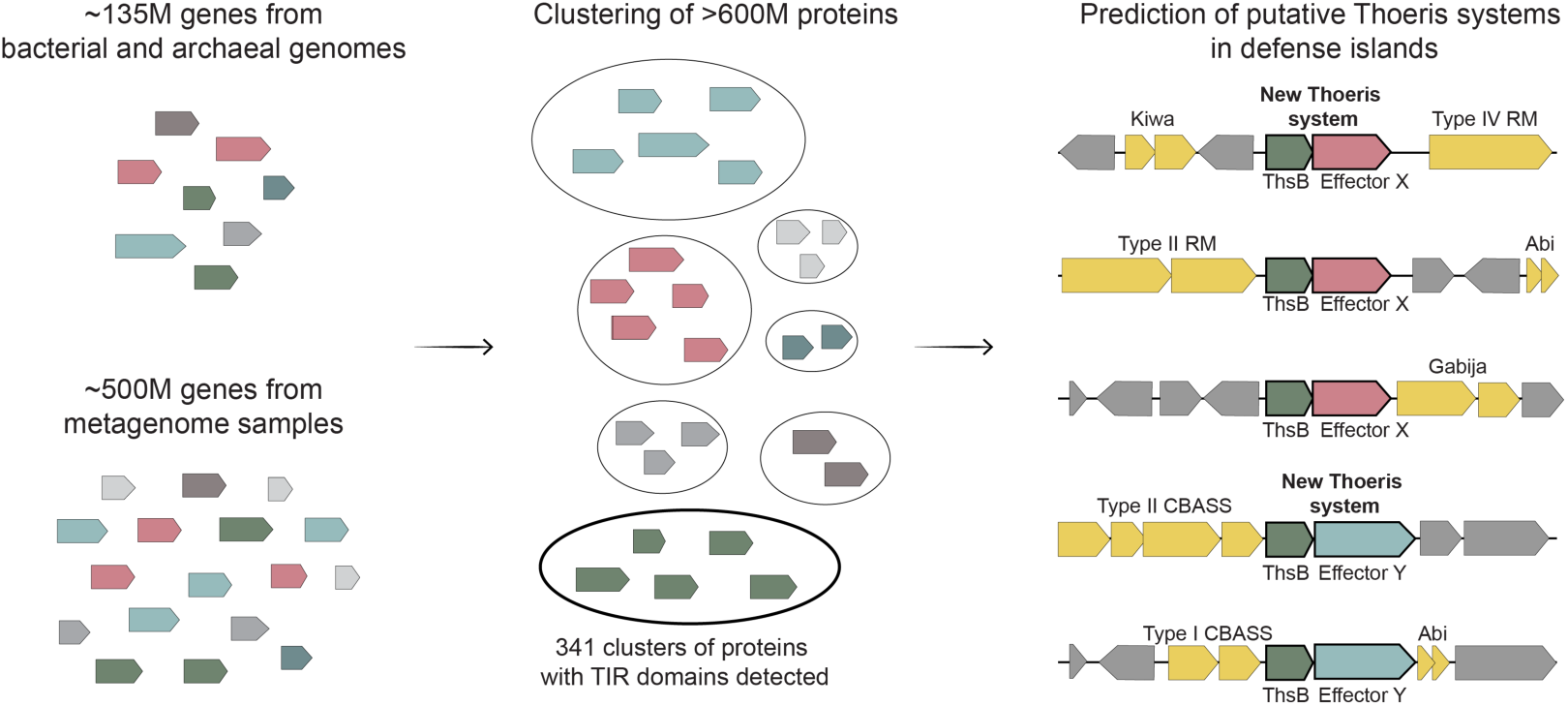
Pipeline for the discovery of novel Thoeris types in microbial genomes and metagenomes. Related to figure 1.

**Figure S2.**
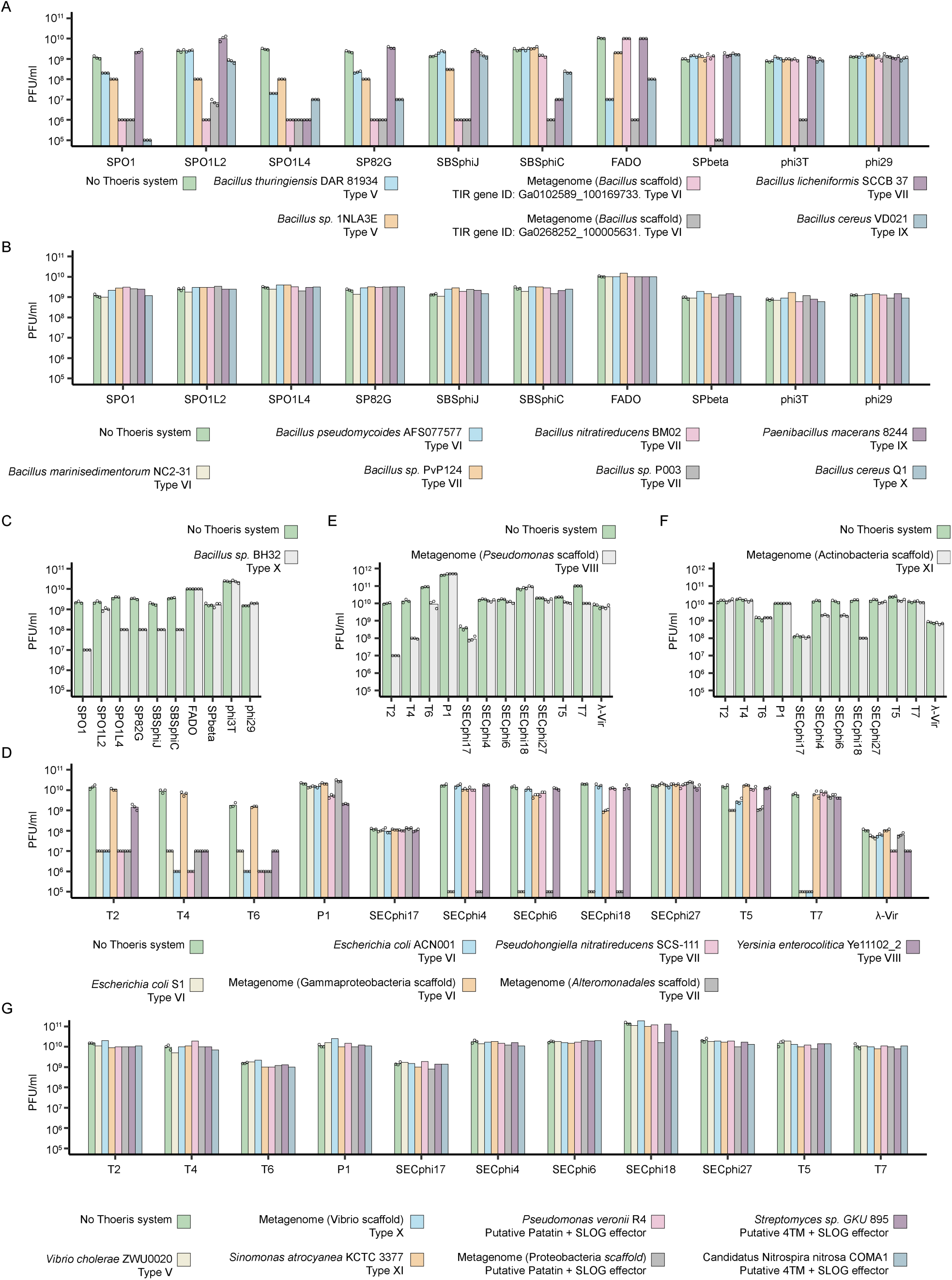
Plaque assay results of experimentally tested Thoeris systems. Phage infection assays showing plaque-forming units (PFU/ml) for control cells and cells expressing indicated Thoeris systems. (A) *B. subtilis* cells expressing the indicated Thoeris systems under the control of their native promoters at room temperature. (B) *B. subtilis* cells expressing candidate systems not verified as defensive, under the control of their native promoters at room temperature. (C) *B. subtilis* cells expressing the type X Thoeris system under a xylose-inducible promoter at room temperature. (D) *E. coli* cells expressing Thoeris systems integrated into the chromosome under the control of an anhydrotetracycline (aTC)-inducible promoter at room temperature. (E) *E. coli* cells expressing the type VIII Thoeris system from a metagenomic scakold under the control of an aTC-inducible promoter at 37 °C. (F) *E. coli* cells expressing the type XI Thoeris system from a plasmid under the control of an arabinose-inducible promoter at room temperature. (G) *E. coli* cells expressing candidate systems not verified as defensive, from plasmids under the control of an arabinose-inducible promoters at room temperature. Bars represent the mean of three replicates for defensive systems, with individual data points overlaid; non-defensive systems were tested once. The same Negative control data (“No Thoeris”) are presented in panels S2A and S2B. The data for the type V Thoeris system from *B. thuringiensis* DAR 81934 presented in panels S2A and 3E are the same. The data for phage SBSphiJ infecting negative control cells and cells expressing the type VI Thoeris system from the metagenomic *Bacillus* scaffold presented in panels S2A and 4D are the same. The data for the type VII Thoeris system from the metagenomic *Alteromonadales* scaffold presented in panels S2D and 5E are the same. The data for the type VIII Thoeris system from the metagenomic *Pseudomonas* scaffold presented in panels S2E and 5F are the same. Related to figure 2.

**Figure S3.**
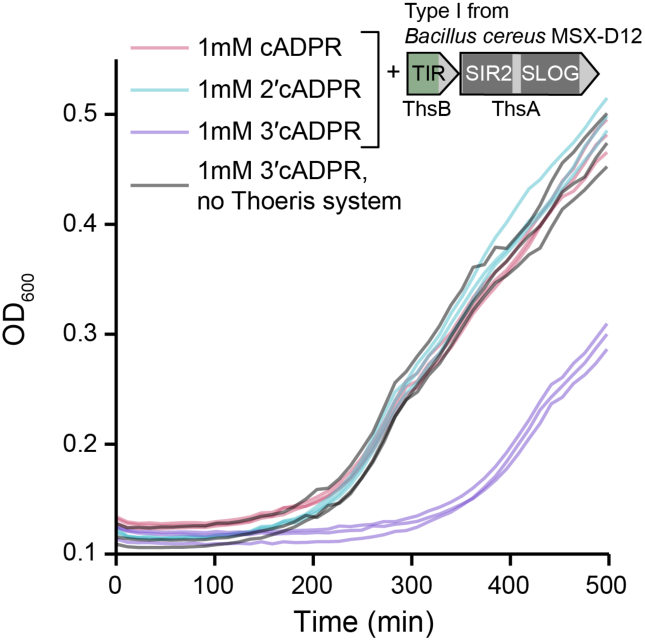
3′cADPR activates ThsA-mediated toxicity in type I Thoeris. Growth curves of *B. subtilis* cells expressing the type I Thoeris system from *Bacillus cereus* MSX-D12 or an empty vector. The growth medium was supplemented with the indicated cyclic ADPR isomers. Three repeats for each condition are shown as individual curves. The data for the “no Thoeris system” control is also presented in panels 3D, S4A and S4B. Related to figure 3.

**Figure S4.**
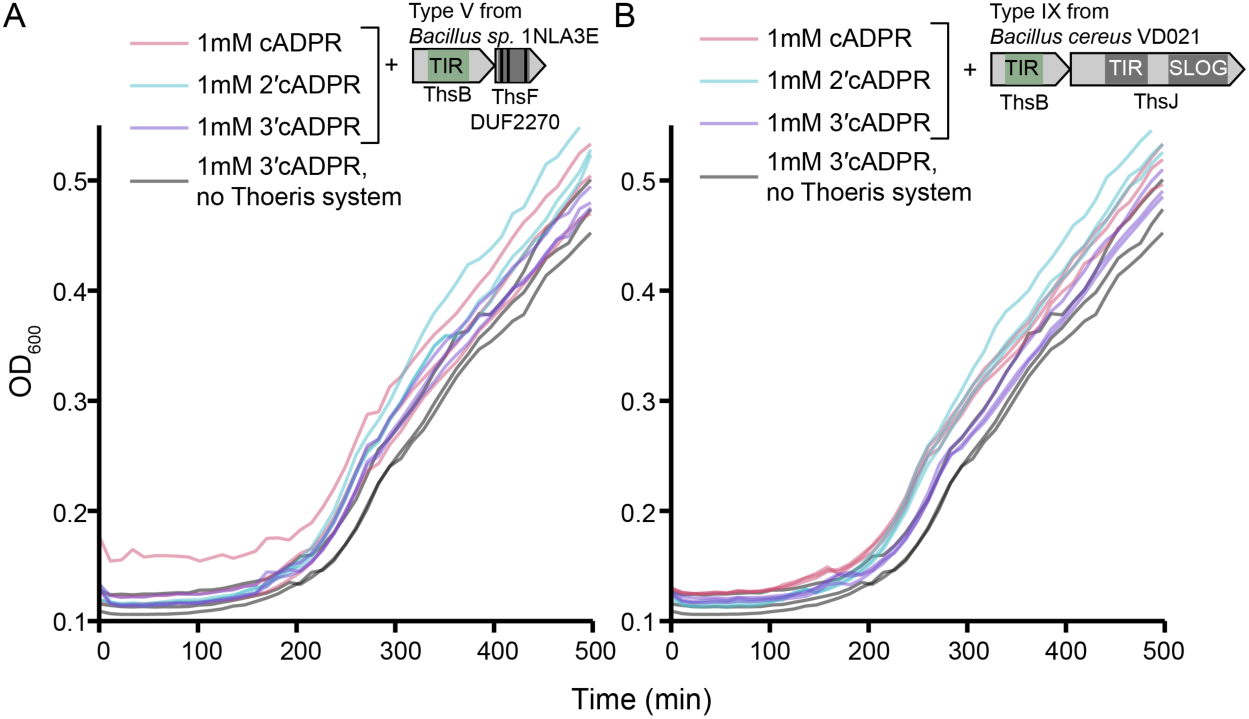
Supplementation of 3′cADPR to the growth medium does not activate eYector-mediated toxicity in the type V Thoeris system from *Bacillus sp.* 1NLA3E and in type IX Thoeris. Growth curves of *B. subtilis* cells expressing the type V Thoeris system from *Bacillus sp.* 1NLA3E (A), the type IX Thoeris system from *Bacillus cereus* VD021 (B), or an empty vector. Growth media were supplemented with the indicated cyclic ADPR isomers. Three repeats for each condition are shown as individual curves. The control “no Thoeris system” data are the same in panels A and B, and are also presented in panels 3D and S3. Related to figure 3.

**Figure S5.**
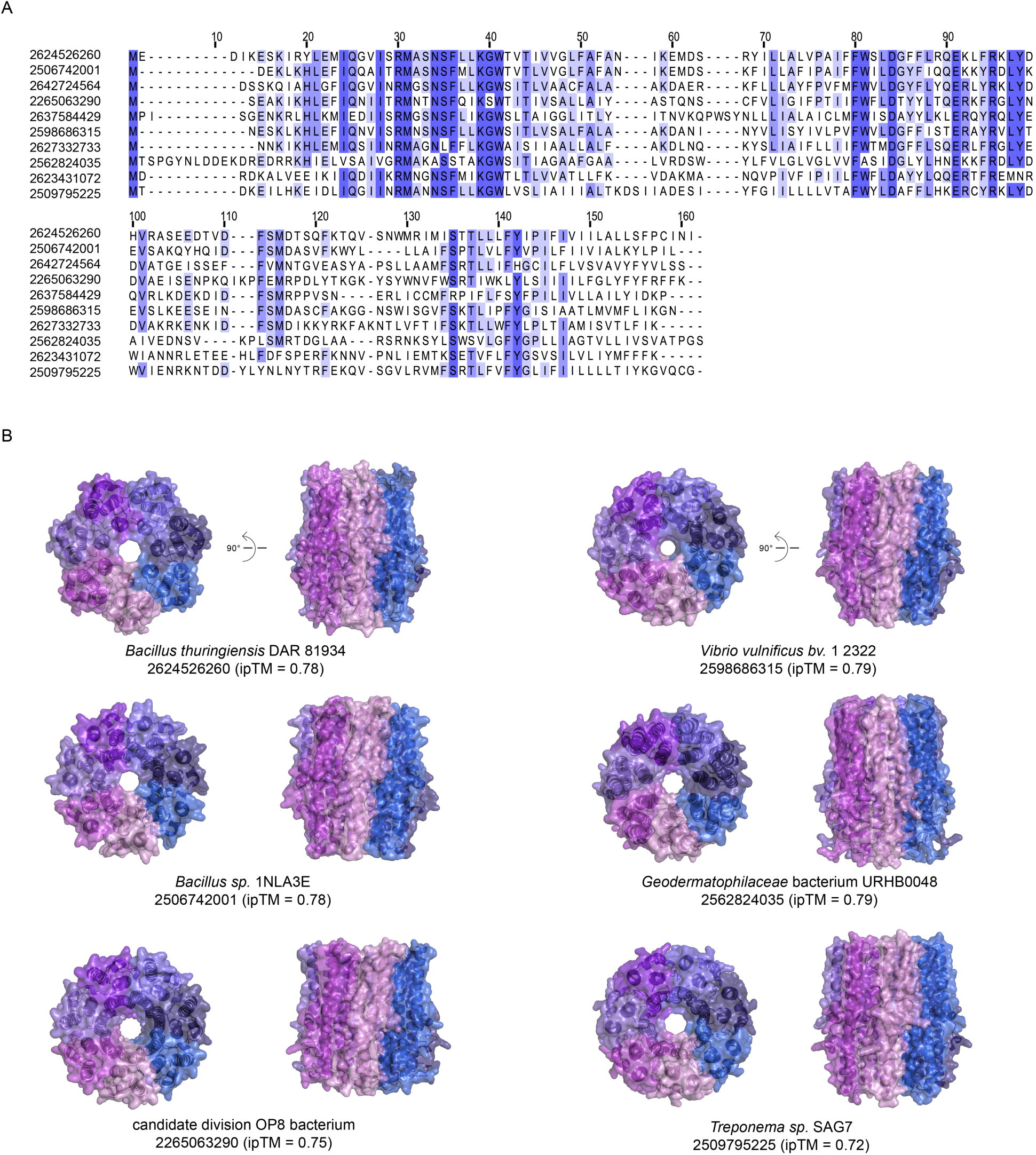
Multiple sequence alignment and structure predictions of ThsF homologs. (A) Multiple sequence alignment of ThsF homologs from type V Thoeris systems, with conserved residues highlighted in purple. IMG gene IDs are shown on the left. (B) Structure predictions of ThsF homologs modeled as homo-heptamers. For each homolog, the organism of origin, IMG gene ID, and ipTM Alphafold3 score of the predicted structure are indicated below. Related to figure 3.

**Figure S6.**
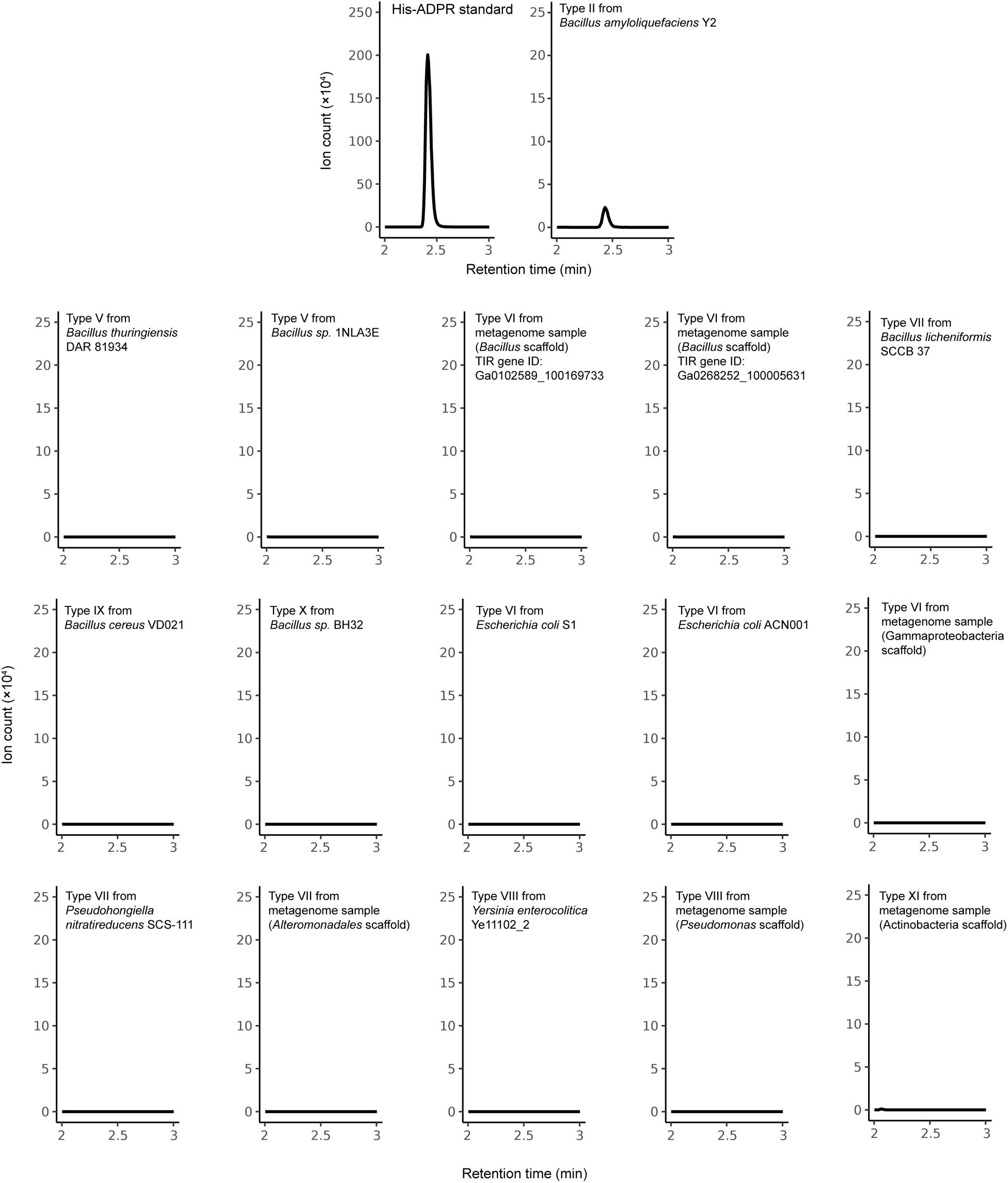
TIR domains from Thoeris types V-XI do not produce detectible His-ADPR in response to infection. Extracted ion chromatograms (m/z = 695.1257) from lysates of cells expressing the TIR domain of the indicated Thoeris system after phage infection. Data on the infecting phage for each strain and the sampling time are provided in Table S2. The chromatogram of the His-ADPR standard (100 µM) is shown above. Data are representative of three replicates. Related to figures 1-5.

**Figure S7.**
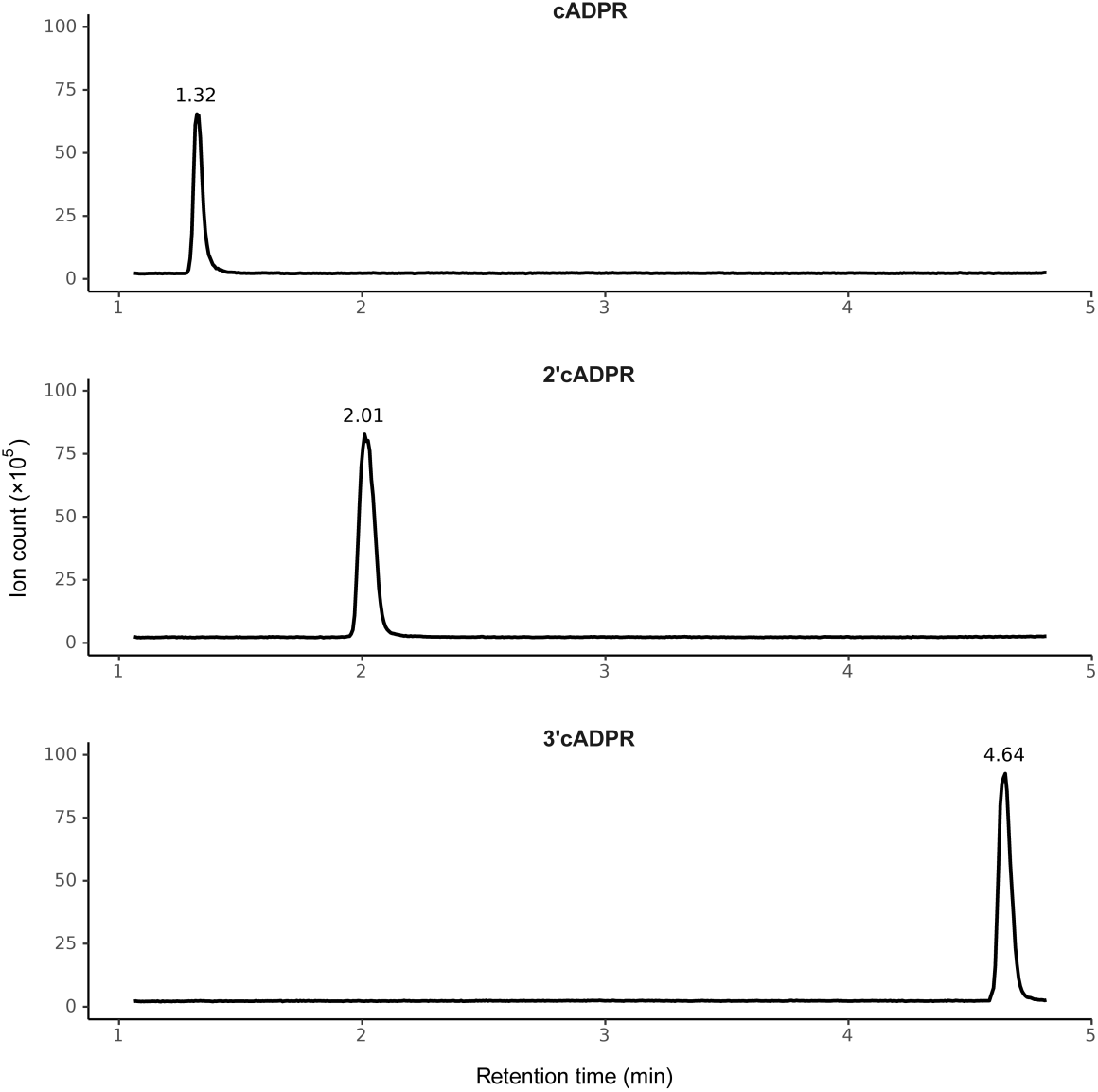
An LC–MS protocol enabling separation of cADPR isomers. Extracted ion chromatograms of cADPR, 2′cADPR, and 3′cADPR standards showing distinct retention times, enabling reliable separation of the three cyclic ADPR isomers. Related to Figures 3–5.

**Figure S8.**
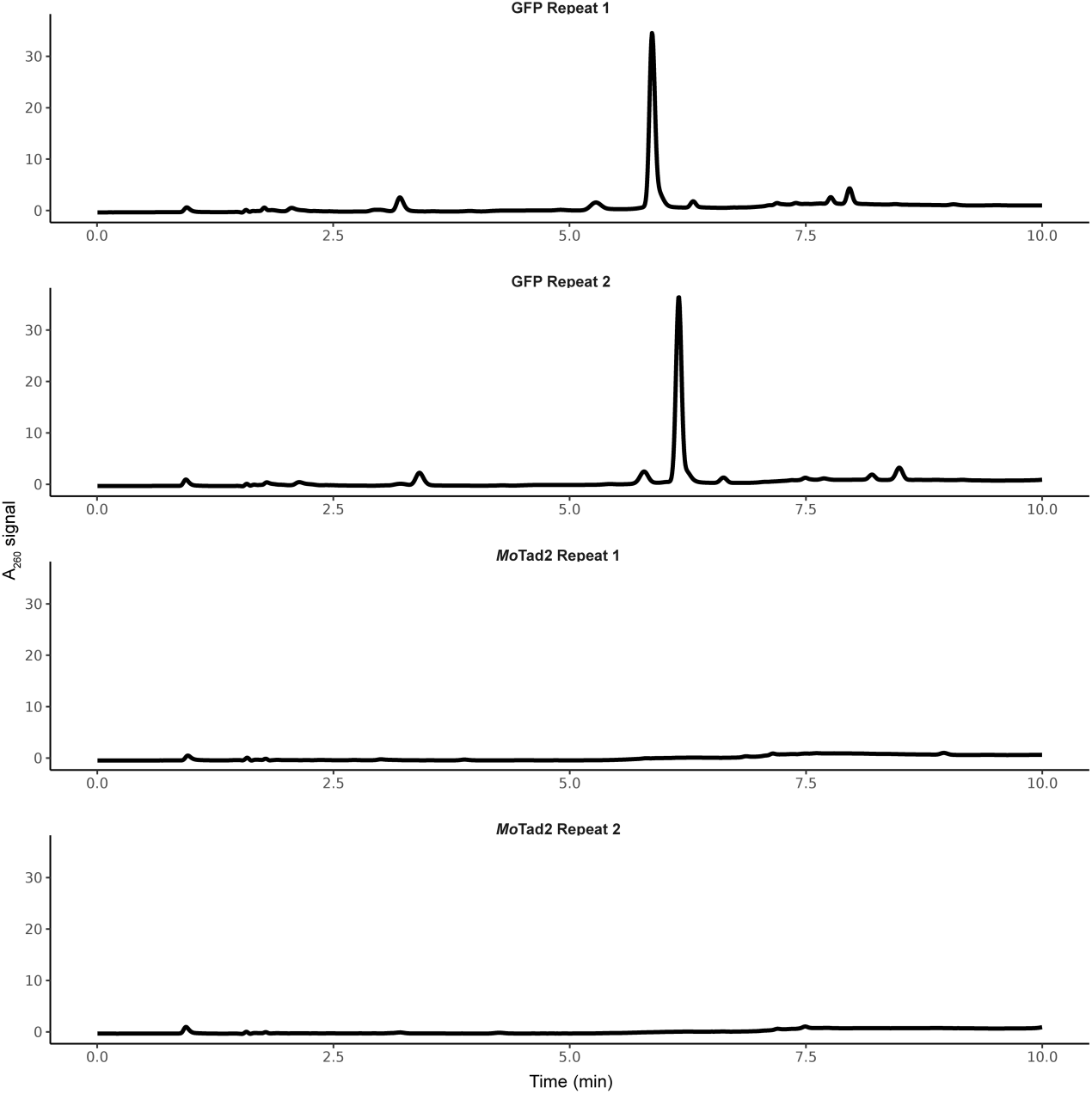
*Mo*Tad2 sequesters 2′cADPR. HPLC analysis of 2′cADPR incubated with purified GFP (control; top lanes) or purified *Mo*Tad2 (bottom lanes) at a 4:1 protein-to-molecule ratio. Related to figure 4.

**Figure S9.**
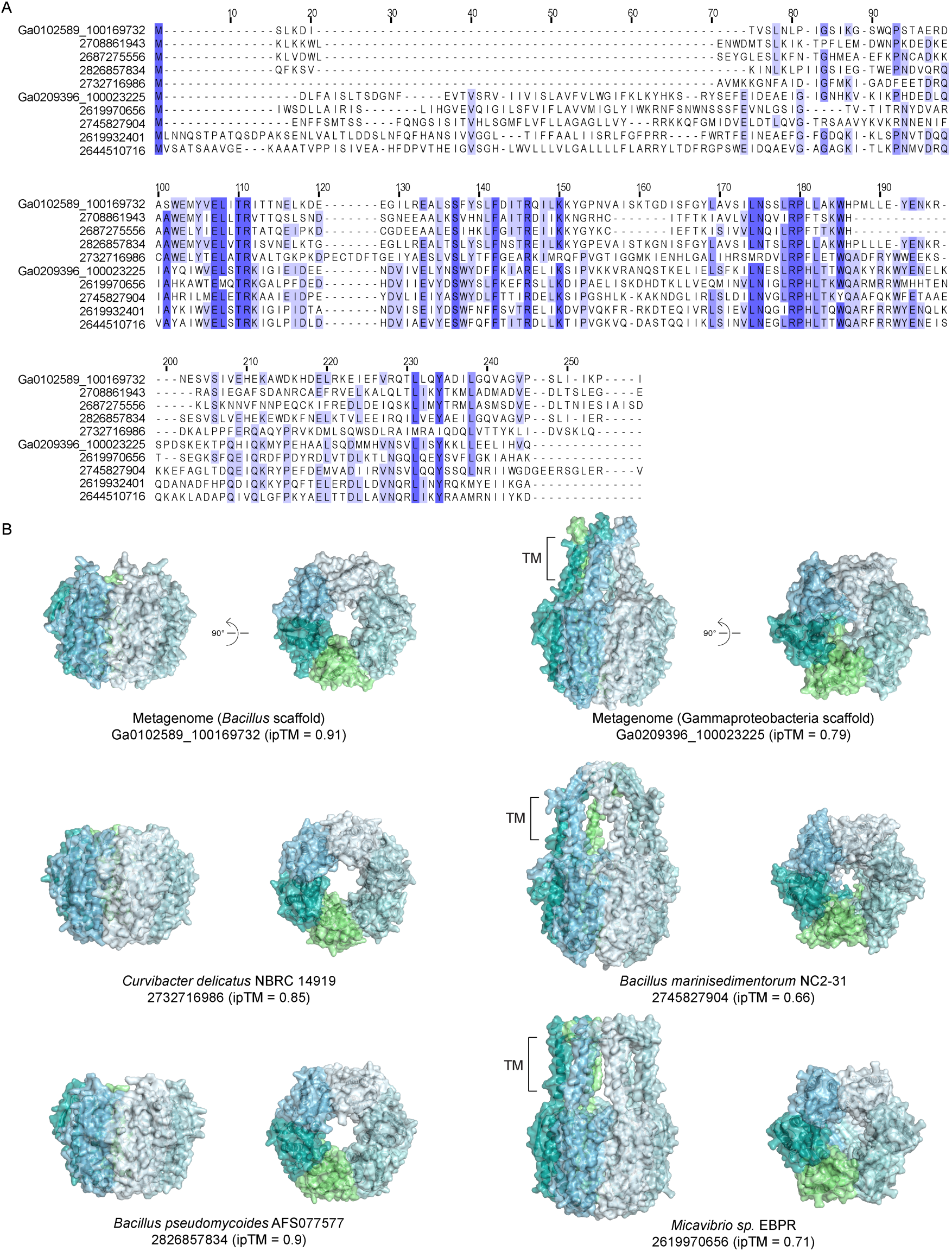
Multiple sequence alignment and structure predictions of ThsG homologs. (A) Multiple sequence alignment of ThsG homologs from type VI Thoeris systems, with conserved residues highlighted in purple. IMG gene IDs are shown on the left. (B) Structure predictions of ThsG homologs modeled as homo-hexamers. For each homolog, the organism of origin, IMG gene ID, and ipTM Alphafold3 score of the predicted structure are indicated below. Related to figure 4.

**Figure S10.**
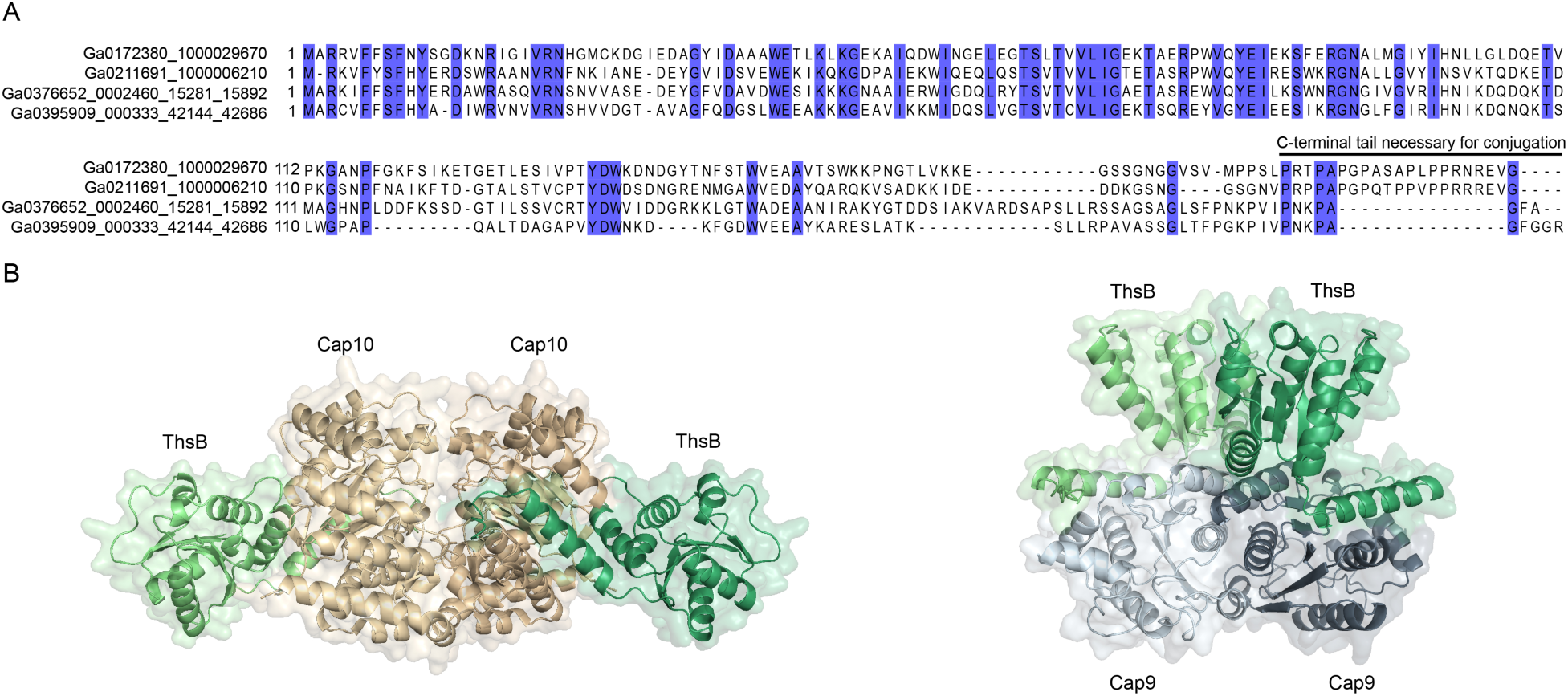
Thoeris operons associated with ancillary proteins typical of CBASS. (**A**) Multiple sequence alignment of ThsB proteins associated with Cap2 and Cap3, with conserved residues highlighted in purple. IMG gene IDs are shown on the left. The C-terminal tail required for Cap2 conjugation, containing the conserved glycine residue, is indicated. (**B**) AlphaFold3 modeling predicts that the ThsB protein from *Pseudomonadales bacterium* SP163 interacts with the corresponding Cap10 (ipTM = 0.79) and Cap9 (ipTM = 0.62) proteins in a manner analogous to the CD-NTase–Cap9/10 interactions observed in type IV CBASS systems. Related to Figure 6.

## Methods

### Bacterial strains

*E. coli* K-12 MG1655 and *B. subtilis* BEST7003 strains were grown in magnesium manganese broth (MMB; LB + 0.1 mM MnCl2 + 5 mM MgCl2) at 37 °C with shaking at 200 RPM. Whenever applicable, the appropriate antibiotics were added at the following concentrations: For *B. subtilis* strains spectinomycin (100 μg mL^−1^) and chloramphenicol (5 μg mL^−1^), and for *E. coli* strains spectinomycin (50 μg mL^−1^) and ampicillin (100 μg mL^−1^).

Type I Thoeris from *Bacillus cereus* MSX-D12 and all other Thoeris systems of *Bacillus* or *Paenibacillus* origin, except for the type X Thoeris from *Bacillus sp.* BH32, were cloned under their native promoters into the *amyE* locus of the *B. subtilis* BEST7003 genome as previously described^5^. A GFP-expressing vector was integrated as a negative control. Type X Thoeris from *Bacillus sp.* BH32 was cloned under a xylose-inducible promoter into the *amyE* locus of *B. subtilis* BEST7003 as previously described^5,6^, with a vector expressing the type I Thoeris ThsB protein from *B. cereus* MSX-D12, under the same xylose-inducible promoter, serving as a negative control for phage infection assays.

Thoeris systems tested in *E. coli* were synthesized and cloned either into the pSG plasmid under their native promoters (type VI systems from *Escherichia coli* S1 and *Escherichia coli* ACN001) or into the pBAD plasmid under an arabinose-inducible promoter (all other systems tested in *E. coli*) and transformed into *E. coli* as previously described^15^. A GFP-expressing plasmid was used as a negative control. An initial plaque assay was performed with these systems, and systems showing defense were then integrated into the *E. coli* MG1655 genome using the Tn7 integration plasmid as previously described^30^ (except for the type XI system from the Actinobacteria scaffold, which was not integrated into the *E. coli* MG1655 genome). These included Thoeris type VI from *Escherichia coli* S1, type VI from *Escherichia coli* ACN001, type VI from the metagenomic Gammaproteobacteria scaffold, type VII from *Pseudohongiella nitratireducens* SCS-111, type VII from the *Alteromonadales* metagenomic scaffold, and both type VIII Thoeris systems. A plasmid expressing RFP was integrated as a negative control.

Anti-defense genes were cloned and transformed as previously described^7,30^. Briefly, Tad1 and *Mo*Tad2 genes were cloned under the control of IPTG promoter into the thrC-Phspank vector. The plasmid was then introduced into *B. subtilis* BEST7003 cells, where the respective defense system was integrated into the *amyE* locus, or into *E. coli* K-12 MG1655, where the defense system was integrated downstream of the gmlS gene.

All plasmids were synthesized by Twist Bioscience or GenScript Corporation. Transformations into *E. coli* were performed using standard electroporation or TSS, and into *B. subtilis* using MC medium, as previously described^5^.

### Phage strains

The *B. subtilis* phages SBSphiJ (GenBank: LT960608.1), Spo1L2 (GenBank: OQ921337.1), SPO1L4 (GenBank: OQ921339.1), SBSphiC (GenBank: LT960610.1), and Fado (GenBank: OM236516.1) were previously isolated by us^5,15,28^. The *B. subtilis* phages phi3T (BGSC: 1L1; GenBank: KY030782.1), SPβ (BGSC: 1L5; GenBank: AF020713.1), SPO1 (BGSC: 1P4; GenBank: NC_011421.1), and SP82G (BGSC: 1P5; GenBank: OM236513) were obtained from the BGSC. The *Bacillus* phage phi29 was acquired from DSMZ (DSM 5546; GenBank: NC_011048.1). SBSphiJ with a Tad1 knockin was generated by us previously^7^. SBSphiJ with a *Mo*Tad2 knockin was generated with the DNA sequence of *Mo*Tad2 under the control of Tad1 promoter as described before^7^, except that the *Mo*Tad2 DNA fragment was synthesized by GenScript Corporation rather than amplified from a phage genome.

The *E. coli* phages T2 (DSMZ 16352) and T6 (DSMZ 4622) were obtained from DSMZ. The *E. coli* phages T5 (GenBank: AY543070.1), λ-vir (GenBank: NC_001416.1), T7 (GenBank: NC_001604.1), T4 (GenBank: AF158101.6), and P1 (GenBank: AF234172.1) were kindly provided by Prof. Udi Qimron. The *E. coli* phages SECphi4 (GenBank: MT331608.1), SECphi6 (GenBank: CADCZA020000001.1), SECphi17 (GenBank: LT960607.1), SECphi18 (GenBank: LT960609.1), and SECphi27 (GenBank: LT961732.1) were previously isolated by us^5,56^.

All phages were propagated on either *E. coli* MG1655 or *B. subtilis* BEST7003 by picking a single phage plaque into a liquid culture grown at 37 °C to an optical density at 600 nm (OD600) of 0.3 in MMB broth until culture collapse (or 3 h in the case of no lysis). The culture was then centrifuged for 10 min at 3,200 *g* and the supernatant was filtered through a 0.2 µm filter to get rid of remaining bacteria and bacterial debris.

### Generating protein clusters

A database of ∼135 million proteins derived from ∼38,000 bacterial and archaeal genomes was downloaded from the Integrated Microbial Genomes (IMG)^57^ database in October 2017, and all protein sequences were aggregated into protein clusters as previously described^15^.

Next, this dataset was further expanded with protein sequences from metagenome samples. For this, all scaffolds containing more than 20 annotated genes from ∼25,000 metagenome samples were downloaded from the IMG database in May 2020, resulting in ∼500 million coding genes. The protein sequences from these scaffolds were filtered for redundancy using the “clusthash” option of MMseqs2^58^ (release 13-45111) with the parameter “--min-seq-id 0.9”, yielding a set of ∼220 million unique metagenome protein sequences.

The metagenome protein sequences were then searched against the representative sequences of the isolate-derived clusters using the “search” option of MMseqs2 with the parameters “--max-accept 100 -s 7.5”. Metagenome sequences were added to the closest isolate cluster if a significant similarity was detected (e-value < 0.001). Metagenome protein sequences that did not match any isolate clusters were clustered with MMseqs2 using the same pipeline applied in Millman et al.^15^ to generate metagenome-exclusive clusters.

For each resulting protein cluster, the fraction of proteins that were genomically associated with known defense genes (“defense score”) was then calculated as previously described^15^.

### Identification of TIR protein clusters

Pfam^59^ annotations for all proteins in the dataset were downloaded from the IMG database. Protein clusters were annotated as TIR clusters if the most common Pfam annotation among proteins in the cluster corresponded to one of the following Pfams: PF18567, PF01582, PF08937, PF10137, PF13676, or PF08357, and at least 20% of the proteins in the cluster carried that annotation.

Additionally, multiple sequence alignments were generated for each cluster using Clustal Omega^60^ version 1.2.4, and the alignments were searched using the “hhsearch” option of hhsuite^61^ version 3.0.3 against the pfam_v32 database, with the parameters “-p 10 -loc -z 1 - b 1 -ssm 2 -sc 1 -seq 1 -dbstrlen 10000 -maxres 32000 -M 60”. Clusters with a hit probability above 90% to any of the Pfams listed above were also defined as TIR clusters.

### Detection of putative Thoeris systems

Multi-gene defense systems were predicted as previously described^5,15^. A total of 341 TIR clusters with a defense score higher than 0.4 and containing at least 10 genes located on genomic scaffolds of sufficient length (having at least 10 flanking genes on each side of the anchor TIR domain protein) were collected. The predicted systems associated with these TIR clusters were examined manually, and putative Thoeris systems were identified. Multi-gene systems in which one of the proteins was annotated with the pfam annotations PF01909 (CBASS^18,19^) or PF00211 (Pycsar^20^), as well as multi-gene systems in which one of the proteins had an hhpred^61^ hit to prokaryotic Argonautes^21,22^ with probability above 90% were discarded. Candidate Thoeris systems were then manually selected for experimental verification to represent both cluster diversity and effector diversity.

### Extraction of TIR domain sequences from putative Thoeris systems

All TIR domain-containing proteins were collected from putative Thoeris systems, along with a set of effector TIR proteins from CBASS and Pycsar systems that were added for reference. A structural model for each protein was generated using AlphaFold3^62^ with default parameters. Each structure was then searched against a curated set of well-defined TIR domain structures (Table S4) using Foldseek^63^ version 5.53465f0. TIR domain boundaries within each protein were defined by identifying the residues corresponding to the median N-terminal and C-terminal positions of hits to the reference TIR domain structures.

### Structural phylogenetic analysis of TIR domains

TIR domain sequences were clustered using the MMseqs2^58^ “cluster” option with the parameters “-c 0.8 --min-seq-id 0.8” to remove redundancy, and a representative TIR domain was selected from each group of homologs.

TIR domain representatives were aligned using FoldMason^64^ version 2.7bd21ed to generate a structure-guided multiple sequence alignment. A phylogenetic tree was subsequently constructed using IQ-TREE^65^ version 2.2.0 with the parameters “-B 1000 -T AUTO” using the default amino acid substitution model. The resulting tree was visualized with iTOL^66^ v6.

The taxonomic source of metagenomic scaffolds encoding Thoeris systems was inferred by extracting the top BLAST^67^ nr hits for all genes located on each scaffold harboring a Thoeris system. For each gene, the organism associated with its top hit was recorded, and the taxonomic origin of the scaffold was assigned based on the majority consensus of these annotations.

### Plaque assays

Phage titer was determined using the small drop plaque assay method^68^. An overnight culture of *E. coli* (300 µL) or *B. subtilis* (400 µL) was mixed with 30 mL MMB 0.5% agar, poured on 10 cm square plates and left to dry for 1 h at room temperature.

For *E. coli* systems integrated into the genome, anhydrotetracycline (aTc) was added to the growth medium at the following concentrations: 10 nM (type VI systems from *Escherichia coli* S1 and *Escherichia coli* ACN001), 100 nM (type VI system from the metagenomic Gammaproteobacteria scaffold, type VII from the *Alteromonadales* metagenomic scaffold, type VIII from the metagenomic *Pseudomonas* scaffold, and the negative control), and 200 nM (type VII from *Pseudohongiella nitratireducens* SCS-111 and type VIII from *Yersinia enterocolitica* Ye11102_2). For *E. coli* systems harboring constructs under the arabinose-inducible pBAD promoter, 0.2% arabinose was added to the medium. Expression of the type X Thoeris system from *Bacillus sp.* BH32 was induced with 0.2% xylose. In bacterial strains expressing Tad1 or *Mo*Tad2, the medium was supplemented with 1mM IPTG.

Tenfold serial dilutions in MMB were carried out for each of the tested phages and 10 µL drops were put on the bacterial layer. After the drops had dried up, the plates were inverted and incubated overnight at room temperature (except for the type VIII Thoeris system from the metagenomic *Pseudomonas* scaffold, which was incubated at 37 °C). PFUs were determined by counting the derived plaques after overnight incubation. When no individual plaques could be identified, a faint lysis zone across the drop area was considered to be 10 plaques. The efficiency of plating was measured by comparing plaque assay results for control bacteria and those for bacteria containing the respective Thoeris system.

### Cell lysates preparation for characterization of ThsB-derived molecules

*E. coli* MG1655 cells carrying a plasmid expressing ThsB from the new Thoeris systems were grown at 25 °C (except for cells expressing ThsB from the type VIII Thoeris system from the metagenomic *Pseudomonas* scaffold, which were grown at 37 °C) with shaking at 200 rpm until reaching an OD₆₀₀ of 0.3 in 150 mL MMB medium.

ThsB proteins of the type VI Thoeris system from the metagenomic Gammaproteobacteria scaffold, type VII from *Pseudohongiella nitratireducens* SCS-111, type VII from the Alteromonadales metagenomic scaffold, both type VIII systems, and the type XI system from the metagenomic Actinobacteria scaffold were cloned into the pBAD plasmid under the arabinose-inducible promoter, whereas the type VI systems from *Escherichia coli* S1 and *Escherichia coli* ACN001 were cloned into the pSG plasmid under their native promoters.

All ThsB proteins with a *Bacillus* or *Paenibacillus* origin were cloned under their native promoters into the *amyE* locus of the *B. subtilis* BEST7003 genome, except for the type X Thoeris system from *Bacillus* sp. BH32, which was cloned under a xylose-inducible promoter into the *amyE* locus of *B. subtilis* BEST7003.

Cells were then infected with phage as indicated in Table S2 at an MOI of 3 for *E. coli* or an MOI of 10 for *Bacillus*. Samples were collected before and after infection at the timepoints listed in Table S2. A total of 50 mL of cells was centrifuged at 4 °C, 4,000 *g* for 10 min, and the pellet was frozen in liquid nitrogen and stored at −80 °C.

To extract cell metabolites, 600 μL of 100 mM sodium phosphate buffer (pH 7.5) was added to each pellet. Samples were transferred to FastPrep Lysing Matrix B tubes (2 mL; MP Biomedicals, cat. #116911100) and lysed at 4 °C using a FastPrep bead beater for two rounds of 40 s at 6 m s^−1^. Tubes were then centrifuged at 4 °C for 10 min at 15,000 *g*. The supernatant was transferred to an Amicon Ultra-0.5 3 kDa centrifugal filter unit (Merck Millipore, cat. no. UFC500396) and centrifuged for 45 min at 4 °C, 12,000 *g*.

Filtered lysates were analyzed by LC-MS, ThsA NADase assays, and ThsE protease reporter assays as described below.

### ThsA NADase assay

ThsA NADase assays were performed as previously described^6,7,28^, with minor modifications. Briefly, in a total volume of 5 µL purified ThsA protein at a final concentration of 100 nM was mixed with 1,N6-ethenoadenine dinucleotide (εNAD; Sigma, N2630) at a final concentration of 0.5 mM and combined with 5 µL of lysates in a black 384-well plate (Corning 4511). Fluorescence was measured on a Tecan Infinite M200 plate reader every minute during incubation at 25 °C, using 300 nm excitation and 410 nm emission wavelengths. The reaction rate was calculated from the linear part of the initial reaction.

### ThsE protease reporter assay

ThsE caspase-like protease, synthetic EEKAR–aminomethylcoumarin (AMC) conjugated peptides, and N7-cADPR were produced as described previously^10^. Five microliters of filtered lysates were mixed with 1 µM ThsE-containing lysate, 2 µL of 100 µM AMC peptides, and 2 µL of 100 mM sodium phosphate buffer (pH 7.4) in a black 384-well plate (Corning 4511). AMC fluorescence (excitation/emission: 341/441 nm) was measured on a Tecan Infinite M200 plate reader at 37 °C every 2 minutes and quantified relative to a 10 µM AMC standard solution.

### LC-MS analysis of filtered lysates

The sample solutions were analyzed by UPLC–HRMS without dilution. The analyses were carried out on a Waters SYNAPT-XS Q-Tof mass spectrometer (Manchester, UK) with an electrospray ionization (ESI) source.

The spectra were recorded in the negative ion mode within a mass range from 100 to 1200 m/z. The parameters were set as follows: capillary voltage at 1.5 kV, cone gas flow at 50 L/h, source temperature at 140°C, and cone voltage at 20 V (−). The desolvation temperature was set at 600°C, and the desolvation gas (N2) flow rate was set at 800 L/h.

Lock spray was acquired with Leucine Encephalin (m/z=554.2615 in negative mode) at a concentration of 200 ng/mL and a flow rate of 10 μL/min once every 10 s for a 1 s period to ensure mass accuracy. Waters MassLynx v4.2 software was used for data acquisition and data processing. Standard molecules (2′cADPR, 3′cADPR, cADPR, and His-ADPR) were provided by BIOLOG and analyzed in the same conditions as references. Signals with m/z of cADPR isomers (540.053) and His-ADPR (695.126) were extracted with ±5 ppm mass accuracy, and their retention times were compared with the standards.

The analytes were separated using the Waters Premier Acquity UPLC system. The gradient elution was achieved with a Waters Acquity Premier HSS T3 Column, 1.8 μm, 2.1 × 100 mm at 0.4 mL/min flow rate, 35°C. Mobile phase A consisted of 20 mM aqueous Ammonium Acetate (Sigma-Aldrich 09691) and mobile phase B consisted of 20 mM Ammonium Acetate in acetonitrile:water ration of 75:25. The gradient was as follows: 0–2.5 min 0% of B, 2.5-6 min 10% of B.

### Molecule-induced toxicity experiments

Overnight bacterial cultures were diluted 1:50 into MMB medium (except for experiments with type VI systems, for which a defined MC medium was used), supplemented or not with 1 mM cADPR, 2′cADPR, or 3′cADPR, and grown in 20 µL volumes in 384-well plates at 25 °C (except for the experiment with the type VIII Thoeris system from the metagenomic *Pseudomonas* scaffold, which was performed at 37 °C). MC medium was composed of 80 mM K2HPO4, 30 mM KH2PO4, 2% glucose, 30 mM trisodium citrate, 22 μg/mL ferric ammonium citrate, 0.1% casein hydrolysate (CAA), 0.2% potassium glutamate. Optical density at 600 nm (OD₆₀₀) was measured every 10 minutes using a Tecan Infinite 200 plate reader.

### Structural analysis of Thoeris proteins

Structural models of Thoeris proteins were generated using AlphaFold3^62^ with default parameters. For Thoeris effectors, homo-oligomeric assemblies were inferred by modeling each effector as a homo-oligomer (between a dimer and an octamer) and selecting the assembly with the highest ipTM score as the predicted oligomeric state. Structural figures were prepared in PyMOL v2.5.4. Electrostatic surface maps were calculated using APBS^69^ v3.1.1 with the default settings implemented in the PyMOL APBS plugin. Multiple sequence alignments of Thoeris effectors presented in figures S5 and S9 were generated using MAFFT^70^ v7.520 with default parameters.

### Detection of Thoeris systems in genomes using DefenseFinder

To generate hidden Markov models (HMMs) for use in DefenseFinder^40^, we first collected, for each Thoeris effector, all genes from its corresponding protein cluster that were located adjacent to *thsB* on the genome, ensuring they originated from *bona fide* Thoeris systems. HMM profiles were constructed as described previously^40^. Briefly, protein sequences were aligned using MAFFT^70^ v7.520 with default parameters, and the resulting alignments were used to generate HMM profiles with hmmbuild (default parameters) from the HMMER^71^ v3.3.2 suite. GA score thresholds for each HMM were set manually based on inspection of score distributions. DefenseFinder definitions for the newly identified Thoeris systems were based on the existing definitions for Thoeris types I–IV.

### *In vitro Mo*Tad2-2′cADPR binding experiment

The *Mo*Tad2 protein used in this study was expressed and purified as previously described^9^. Purified *Mo*Tad2 or GPF were mixed with 2′cADPR in a 4:1 protein-to-molecule ratio to a final concentration of 20 uM, in 100 mM sodium phosphate buffer. Samples were incubated for 10 minutes at room temperature. Following the incubation the samples were filtered through a 3 kDa MWCO filter, and flow through was collected and analyzed by HPLC.

20 µL of the samples were analyzed using HPLC. HPLC of the obtained fraction was performed using Agilent 1260 and a chromatography SUPELCOSIL™ LC-18-T HPLC Column. The following protocol was used for all runs: 1 min of mobile phase A 100%, 2 min 75% A and 25% B, 2 min 50% A and 50% B, 2 min 20% A and 80% B and 3 min 100 % A, 1 mL/min flow rate. Mobile phase A was 20 mM potassium phosphate pH 6 and B was 20 mM potassium phosphate pH 6 in 20% methanol.

## References

1. Essuman, K., Milbrandt, J., Dangl, J. L. & Nishimura, M. T. Shared TIR enzymatic functions regulate cell death and immunity across the tree of life. Science 377, eabo0001 (2022).

2. Fitzgerald, K. A. & Kagan, J. C. Toll-like Receptors and the Control of Immunity. Cell 180, 1044–1066 (2020).

3. Kawasaki, T. & Kawai, T. Toll-Like Receptor Signaling Pathways. Front. Immunol. 5, 461 (2014).

4. Li, S., Manik, M. K., Shi, Y., Kobe, B. & Ve, T. Toll/interleukin-1 receptor domains in bacterial and plant immunity. Current Opinion in Microbiology 74, 102316 (2023).

5. Doron, S. et al. Systematic discovery of antiphage defense systems in the microbial pangenome. Science 359, eaar4120 (2018).

6. Ofir, G., et al. Antiviral activity of bacterial TIR domains via immune signalling molecules. Nature 600, 116–120 (2021).

7. Leavitt, A. et al. Viruses inhibit TIR gcADPR signalling to overcome bacterial defence. Nature 611, 326–331 (2022).

8. Manik, M. K. et al. Cyclic ADP ribose isomers: Production, chemical structures, and immune signaling. Science 377, eadc8969 (2022).

9. Sabonis, D. et al. TIR domains produce histidine-ADPR as an immune signal in bacteria. Nature 642, 467–473 (2025).

10. Rousset, F. et al. TIR signaling activates caspase-like immunity in bacteria. Science 387, 510–516 (2025).

11. Bayless, A. M. et al. Plant and prokaryotic TIR domains generate distinct cyclic ADPR NADase products. Science Advances 9, eade8487 (2023).

12. Wu, Y. et al. A canonical protein complex controls immune homeostasis and multipathogen resistance. Science 386, 1405–1412 (2024).

13. Berg, D. F. van den et al. Bacterial homologs of innate eukaryotic antiviral defenses with anti-phage activity highlight shared evolutionary roots of viral defenses. Cell Host & Microbe 32, 1427–1443.e8 (2024).

14. Wang, S. et al. The role of TIR domain-containing proteins in bacterial defense against phages. Nat Commun 15, 7384 (2024).

15. Millman, A. et al. An expanded arsenal of immune systems that protect bacteria from phages. Cell Host & Microbe 30, 1556–1569.e5 (2022).

16. Hochhauser, D., Millman, A. & Sorek, R. The defense island repertoire of the Escherichia coli pan-genome. PLOS Genetics 19, e1010694 (2023).

17. Gao, L. et al. Diverse enzymatic activities mediate antiviral immunity in prokaryotes. Science 369, 1077–1084 (2020).

18. Cohen, D. et al. Cyclic GMP–AMP signalling protects bacteria against viral infection. Nature 574, 691–695 (2019).

19. Millman, A., Melamed, S., Amitai, G. & Sorek, R. Diversity and classification of cyclic-oligonucleotide-based anti-phage signalling systems. Nat Microbiol 5, 1608–1615 (2020).

20. Tal, N. et al. Cyclic CMP and cyclic UMP mediate bacterial immunity against phages. Cell 184, 5728–5739.e16 (2021).

21. Ni, D., Lu, X., Stahlberg, H. & Ekundayo, B. Activation mechanism of a short argonaute-TIR prokaryotic immune system. Science Advances 9, eadh9002 (2023).

22. Koopal, B. et al. Short prokaryotic Argonaute systems trigger cell death upon detection of invading DNA. Cell 185, 1471–1486.e19 (2022).

22. Gao, L. A. et al. Prokaryotic innate immunity via pattern recognition of conserved viral proteins. Science 377, eabm4096 (2022).

24. Kibby, E. M. et al. Bacterial NLR-related proteins protect against phage. Cell 186, 2410–2424.e18 (2023).

25. Rousset, F. & Sorek, R. The evolutionary success of regulated cell death in bacterial immunity. Current Opinion in Microbiology 74, 102312 (2023).

26. Whiteley, A. T. et al. Bacterial cGAS-like enzymes synthesize diverse nucleotide signals. Nature 567, 194–199 (2019).

27. Hobbs, S. J. & Kranzusch, P. J. Nucleotide Immune Signaling in CBASS, Pycsar, Thoeris, and CRISPR Antiphage Defense. Annual Review of Microbiology 78, 255–276 (2024).

28. Yirmiya, E. et al. Phages overcome bacterial immunity via diverse anti-defence proteins. Nature 625, 352–359 (2024).

29. Hadary, R. et al. Functional diversity of phage sponge proteins that sequester host immune signals. Preprint at 10.1101/2025.08.24.671296 (2025).

30. Tal, N. et al. Structural modeling reveals viral proteins that manipulate host immune signaling. Preprint at 10.1101/2025.07.12.664507 (2025).

31. Yu, H. et al. Activation of a helper NLR by plant and bacterial TIR immune signaling. Science 386, 1413–1420 (2024).

32. Glaría, E. & Valledor, A. F. Roles of CD38 in the Immune Response to Infection. Cells 9, 228 (2020).

33. Partida-Sánchez, S. et al. Cyclic ADP-ribose production by CD38 regulates intracellular calcium release, extracellular calcium influx and chemotaxis in neutrophils and is required for bacterial clearance in vivo. Nat Med 7, 1209–1216 (2001).

34. Partida-Sánchez, S., Randall, T. D. & Lund, F. E. Innate immunity is regulated by CD38, an ecto-enzyme with ADP-ribosyl cyclase activity. Microbes and Infection 5, 49–58 (2003).

35. Duncan-Lowey, B., McNamara-Bordewick, N. K., Tal, N., Sorek, R. & Kranzusch, P. J. Effector-mediated membrane disruption controls cell death in CBASS antiphage defense. Molecular Cell 81, 5039–5051.e5 (2021).

36. Sikowitz, M. D., Cooper, L. E., Begley, T. P., Kaminski, P. A. & Ealick, S. E. Reversal of the Substrate Specificity of CMP N-Glycosidase to dCMP. Biochemistry 52, 4037–4047 (2013).

37. Wagner, A. G. et al. Transition State Analogs of Human DNPH1 Reveal Two Electrophile Migration Mechanisms. J. Med. Chem. 68, 3653–3672 (2025).

38. Hochhauser, D. & Sorek, R. Manipulation of the nucleotide pool in human, bacterial and plant immunity. Nat Rev Immunol (2025) doi:10.1038/s41577-025-01206-w.

39. Burroughs, A. M. & Aravind, L. Identification of Uncharacterized Components of Prokaryotic Immune Systems and Their Diverse Eukaryotic Reformulations. J Bacteriol 202, e00365–20 (2020).

40. Tesson, F. et al. Systematic and quantitative view of the antiviral arsenal of prokaryotes. Nat Commun 13, 2561 (2022).

41. Ledvina, H. E. et al. An E1–E2 fusion protein primes antiviral immune signalling in bacteria. Nature 616, 319–325 (2023).

42. Krüger, L. et al. Reversible conjugation of a CBASS nucleotide cyclase regulates bacterial immune response to phage infection. Nat Microbiol 9, 1579–1592 (2024).

43. Ye, Q. et al. Mechanistic basis for protein conjugation in a diverged bacterial ubiquitination pathway. Nat Struct Mol Biol (2025) doi:10.1038/s41594-025-01696-1.

44. Wassarman, D. R. et al. Deazaguanylation is a nucleobase-protein conjugation required for type IV CBASS immunity. Science 389, 1347–1352 (2025).

45. Wu, Z., Tian, L., Liu, X., Zhang, Y. & Li, X. TIR signal promotes interactions between lipase-like proteins and ADR1-L1 receptor and ADR1-L1 oligomerization. Plant Physiol 187, 681–686 (2021).

46. Wan, L. et al. TIR domains of plant immune receptors are NAD^+^-cleaving enzymes that promote cell death. Science 365, 799–803 (2019).

47. Huang, S. et al. Identification and receptor mechanism of TIR-catalyzed small molecules in plant immunity. Science 377, eabq3297 (2022).

48. Wein, T. & Sorek, R. Bacterial origins of human cell-autonomous innate immune mechanisms. Nat Rev Immunol 22, 629–638 (2022).

49. Wei, W., Graeff, R. & Yue, J. Roles and mechanisms of the CD38/cyclic adenosine diphosphate ribose/Ca2^+^ signaling pathway. World J Biol Chem 5, 58–67 (2014).

50. Fliegert, R., Gasser, A. & Guse, A. H. Regulation of calcium signalling by adenine-based second messengers. Biochem Soc Trans 35, 109–114 (2007).

51. Trebak, M. & Kinet, J.-P. Calcium signalling in T cells. Nat Rev Immunol 19, 154–169 (2019).

52. Coleman, M. P. & Höke, A. Programmed axon degeneration: from mouse to mechanism to medicine. Nat Rev Neurosci 21, 183–196 (2020).

53. Loring, H. S. & Thompson, P. R. Emergence of SARM1 as a Potential Therapeutic Target for Wallerian-type Diseases. Cell Chemical Biology 27, 1–13 (2020).

54. Essuman, K. et al. The SARM1 Toll/Interleukin-1 Receptor Domain Possesses Intrinsic NAD^+^ Cleavage Activity that Promotes Pathological Axonal Degeneration. Neuron 93, 1334–1343.e5 (2017).

55. Horsefield, S., et al. NAD^+^ cleavage activity by animal and plant TIR domains in cell death pathways. Science 365, 793–799 (2019).

56. Millman, A., et al. Bacterial Retrons Function In Anti-Phage Defense. Cell 183, 1551–1561.e12 (2020).

57. Chen, I.-M. A. et al. The IMG/M data management and analysis system v.7: content updates and new features. Nucleic Acids Res 51, D723–D732 (2023).

58. Steinegger, M. & Söding, J. MMseqs2 enables sensitive protein sequence searching for the analysis of massive data sets. Nat Biotechnol 35, 1026–1028 (2017).

59. Mistry, J. et al. Pfam: The protein families database in 2021. Nucleic Acids Res 49, D412–D419 (2021).

60. Sievers, F. & Higgins, D. G. The Clustal Omega Multiple Alignment Package. in Multiple Sequence Alignment: Methods and Protocols (ed. Katoh, K.) 3–16 (Springer US, New York, NY, 2021).

61. Steinegger, M. et al. HH-suite3 for fast remote homology detection and deep protein annotation. BMC Bioinformatics 20, 473 (2019).

62. Abramson, J. et al. Accurate structure prediction of biomolecular interactions with AlphaFold 3. Nature 630, 493–500 (2024).

63. van Kempen, M. et al. Fast and accurate protein structure search with Foldseek. Nat Biotechnol 42, 243–246 (2024).

64. Gilchrist, C. L. M., Mirdita, M. & Steinegger, M. Multiple Protein Structure Alignment at Scale with FoldMason. Preprint at 10.1101/2024.08.01.606130 (2024).

65. Minh, B. Q. et al. IQ-TREE 2: New Models and Efficient Methods for Phylogenetic Inference in the Genomic Era. Mol Biol Evol 37, 1530–1534 (2020).

66. Letunic, I. & Bork, P. Interactive Tree of Life (iTOL) v6: recent updates to the phylogenetic tree display and annotation tool. Nucleic Acids Res 52, W78–W82 (2024).

67. Camacho, C. et al. BLAST+: architecture and applications. BMC Bioinformatics 10, 421 (2009).

68. Mazzocco, A., Waddell, T. E., Lingohr, E. & Johnson, R. P. Enumeration of Bacteriophages Using the Small Drop Plaque Assay System. in Bacteriophages: Methods and Protocols, *Volume* 1*: Isolation, Characterization, and Interactions* (eds. Clokie, M. R. J. & Kropinski, A. M.) 81–85 (Humana Press, Totowa, NJ, 2009).

69. Jurrus, E. et al. Improvements to the APBS biomolecular solvation software suite. Protein Sci 27, 112–128 (2018).

70. Katoh, K., Misawa, K., Kuma, K. & Miyata, T. MAFFT: a novel method for rapid multiple sequence alignment based on fast Fourier transform. Nucleic Acids Res 30, 3059–3066 (2002).

71. Eddy, S. R. Accelerated Profile HMM Searches. PLOS Computational Biology 7, e1002195 (2011).

